# Distinct feedforward and feedback pathways for cell-type specific attention effects

**DOI:** 10.1101/2022.11.04.515185

**Authors:** Georgios Spyropoulos, Marius Schneider, Jochem van Kempen, Marc Alwin Gieselmann, Alexander Thiele, Martin Vinck

## Abstract

Spatial attention selectively enhances neural responses to visual stimuli. There are two long-standing hypotheses about how top-down feedback enhances sensory responses in areas like V4: First, by amplifying V1-to-V4 feedforward communication via 30-80Hz gamma-coherence. Second, via top-down feedback to V4 supra- and infra-granular layers. To test these hypotheses, we recorded distinct cell-types across macaque V1 and V4 layers. Attention increased both V1-V4 gamma-coherence and V4 spike-rates, yet with distinct laminar and cell-type profiles. Surprisingly, V1 gamma did not engage V4 excitatory neurons, but only Layer-4 fast-spiking interneurons. Similar observations were made in mouse visual-cortex, where feedforward gamma-influences preferentially recruit optogenetically-tagged PV+ and narrow-waveform SSt+ interneurons. By contrast, attention enhanced V4 spike-rates in both excitatory neurons and fast-spiking interneurons, with the strongest and earliest modulation in Layer-2/3, consistent with a feedback influence. These findings reveal distinct feedforward and feedback pathways for the attentional modulation of inter-areal coherence and spike rates, respectively.

## Introduction

Attention reflects the ability to selectively process and respond to behaviorally relevant sensory information. Higher mammals like primates have the capacity to direct their attention towards specific stimuli based on learned cue-reward associations. The modulation of neural stimulus-responses caused by top-down attention is thought to be mediated by cortical feedback from frontal areas (e.g. FEF) towards sensory areas (e.g. V4) (Gregoriou et al., 2014; Noudoost et al., 2010). An outstanding question is the nature of the circuit mechanisms through which top-down feedback selectively enhances the sensory processing of attended sensory stimuli.

To answer this question, three major challenges need to be overcome: First, selective attention modulates distinct aspects of neural activity, and does so in multiple cortical areas simultaneously. One consistent finding is the enhancement of stimulus-evoked firing rates, especially in higher levels of the primate ventral stream (Buffalo et al., 2010). This increase in spike rates can be understood as a gain modulation of firing responses to the attended stimulus (McAdams and Maunsell, 1999). Another main finding is an increase in inter-areal phase-locking in the gamma-frequency (30-80Hz) range between areas (Bosman et al., 2012; Grothe et al., 2012; Ferro et al., 2021a). In order to formulate a complete mechanistic account of attention, these two phenomena must be put together in a coherent way. Second, a profound challenge in the dissection of circuit mechanisms is the interwoven nature of feedback and feedforward connections due to the reciprocal connections between cortical areas. For instance, it is possible that feedback from e.g. frontal areas induces changes in V1 activity that lead to subsequent changes in neural activity in downstream areas like V2 and V4 (Fries, 2015). However, it is also possible that feedback modulates activity in e.g. area V4, which, in turn, modulates activity in area V1. This influence from V4 to V1 may, then, lead to subsequent feedforward changes in area V4, thereby closing a functional loop (Buffalo et al., 2010; Debes and Dragoi, 2023). Crucially, feedforward and feedback connections are organized along cortical layers, which therefore enable the separation of information flow related to feedforward and feedback processing (Vezoli et al., 2021a; Van Kerkoerle et al., 2017). Third, the circuit mechanisms of attention likely rely on specific interactions between GABAergic interneurons and excitatory neurons, which are difficult to target, particularly in primates. Previous work has suggested differences in attentional modulation between these cell types (Mitchell et al., 2007; Kim et al., 2016; Vinck et al., 2013), however it remains to be determined whether they show specific modulations related to inter-areal interactions and feedback/feedforward processing. For instance feedforward projections tend to predominantly target excitatory cells and fast-spiking interneurons in the granular layer, whereas feedback targets excitatory cells and a broader set of different interneuronal classes (Batista-Brito et al., 2018; Vezoli et al., 2021a; Shen et al., 2022).

There are two main competing theories of the mechanisms underlying selective attention that have made specific proposals about the nature of feedforward and feedback interactions mediating attentional modulation. According to one major hypothesis, attentional selection leads, via top-down feedback, to the enhancement of feedforward (FF) inter-areal information transmission via interareal oscillatory synchronization (Fries, 2005; Kreiter, 2006; Fries, 2015). The enhanced effective communication is then thought to induce subsequent increases in neural responses (i.e. firing rates) for attended stimuli downstream (Fries, 2005; Kreiter, 2006; Fries, 2015). A competing hypothesis posits that the main effect of selective attention is to enhance the gain of neural responses for attended stimuli via top-down feedback (FB), with the strongest and earliest effects at higher hierarchical levels closer to behavioral responses (Desimone et al., 1995; Maunsell, 2015; Buffalo et al., 2010). Such gain modulation may depend on e.g. the top-down modulation of specific GABAergic interneurons, like SSt+ or VIP+ interneurons (Zhang et al., 2014). These two hypotheses (rate vs. coherence) are not necessarily exclusive, and it is possible that distinct roles are played by these two mechanisms considering the laminar organization of feedforward and feedback (Vezoli et al., 2021a): For instance, enhanced feedforward processing via gamma-synchronization may primarily lead to changes in neural activity in the granular layer 4 of downstream areas, whereas feedback may primarily modulate neural activity in the extra-granular layers.

A major impediment to test these theories and determine the nature of feedforward-feedback interactions is the lack of simultaneous recordings from multiple areas of the primate cortex at a laminar and cell-type specific resolution. Using laminar probes, we recorded local field potentials (LFPs) and well-isolated single units from areas V1 and V4 in macaque monkeys performing a spatial attention task. We then analyzed the way in which distinct cell classes in area V4, across different cortical laminae, were phase-locked to the V1 gamma-rhythm, and how this long-range phase-locking was modulated by attention. We contrasted these effects with the cell- and layer-specific attentional modulation of stimulus-driven firing rates, and performed population decoding analyses to determine which signals tracked attention reliably from trial-to-trial. Finally, we analyzed Neuropixels data from mice and investigated the inter-areal phase-locking of different GABAergic subtypes, which were identified through optogenetic tagging.

## Results

We recorded laminar Local Field Potentials (LFPs) and single-unit spiking-activity simultaneously from areas V1 and V4, (Fig. 1A), while monkeys performed an attention task (Fig. 1B). Consistent with previous studies in macaques (Bosman et al., 2012; Grothe et al., 2012), visual stimulation increased gamma-band phase-locking between V1 and V4 LFPs (Fig. 1C). Several further analyses indicated that V1-LFP-V4-LFP gamma phase-locking did not result from coupling of intrinsic V1 and V4 oscillations, but from the feedforward propagation of the V1 gamma rhythm: (1) LFP-LFP locking was accompanied by only weak V4-spike-to-V1-LFP locking, especially when compared to local V1-spike-to-V1-LFP locking (Fig. 1D and F); (2) V4-spike-to-V4-LFP locking did not display a clear peak in the gamma range (fig. 1E); and (3) on average, V1 cells spiked at earlier V1 gamma phases than V4 cells (mean phase difference = 1.51 radians) (Fig. 1F and G).

**Fig. 1:**
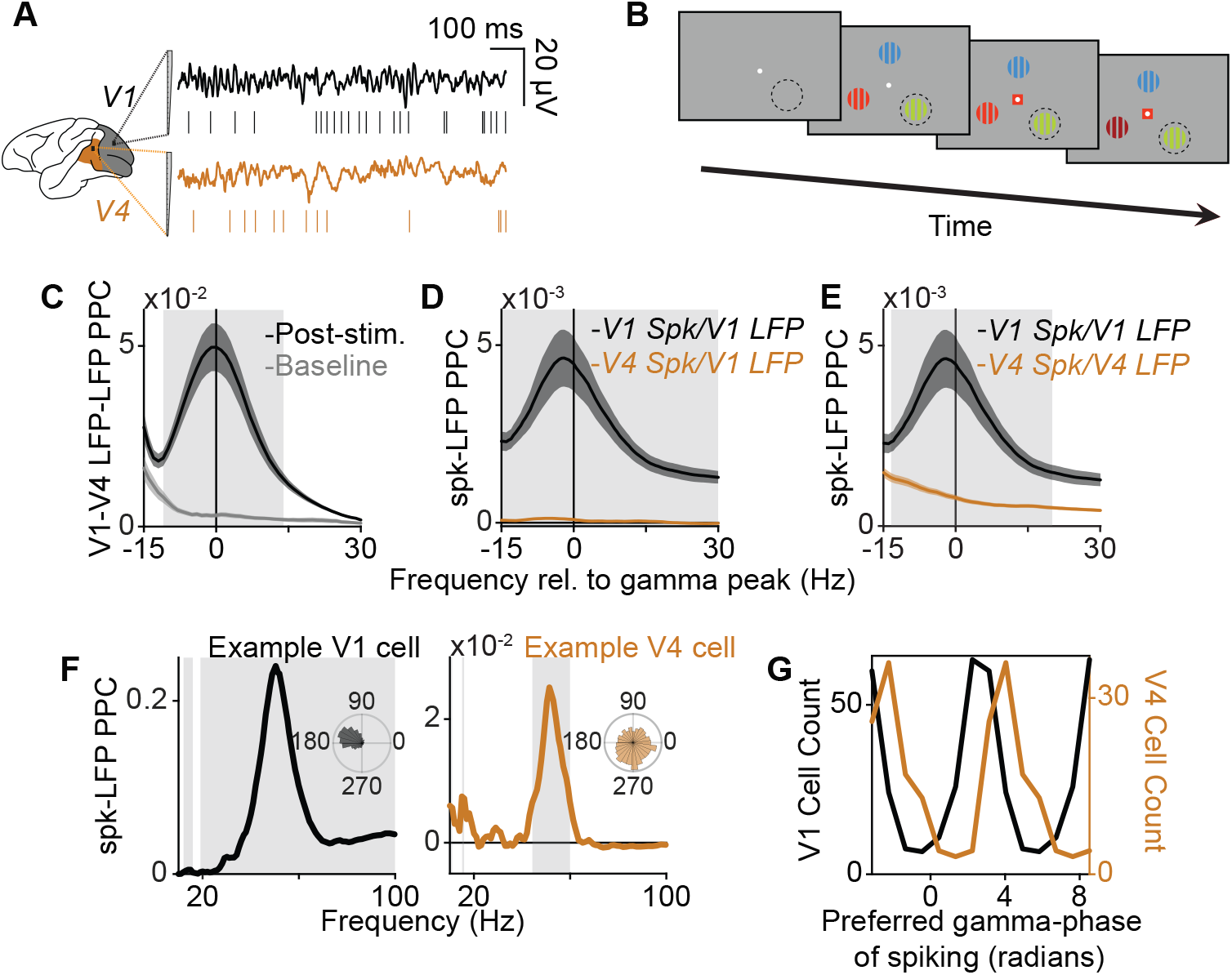
Feedforward gamma-band synchronization between V1 and V4 in the monkey. **(A)** Experimental setup in the macaque. **(B)** Illustration of selective attention task. **(C)** Phase-locking (PPC) between V1 and V4 LFPs during the baseline and visual stimulation period (*N* = 68 sessions). **(D)** PPC between V1 LFPs and V1 single units (SU) (*N* = 311) or V4 SUs (*N* = 397). **(F)** Same as Fig. 1D, but for V1 LFPs and V1 SU spiking, and V4 LFPs and V4 SU spiking. **(G)** Mean V1-gamma phase of spiking for significantly gamma-locked V1 or V4 SUs (mean phase difference = 1.51 radians, *p* = 0, randomization test). **(C-E)** Confidence intervals designate SEM across cells, gray rectangles designate significantly different frequency bins, randomization test between cell-types, FDR correction for multiple comparisons with a threshold of *P* < 0.05). **(F)** Gray rectangles designate significantly different frequency bins, Rayleigh’s test for uniformity, Bonferonni correction for multiple comparisons with a threshold of *P* < 0.05).

To gain deeper physiological insight into these observations, we analyzed the waveform characteristics of V4 single-unit spiking. Units could be clearly separated into two main classes with broad (BW) and narrow (NW) waveforms (Fig. 2A) corresponding to putative excitatory and fast-spiking (FS) interneurons, respectively (McCormick et al. (1985); Mitchell et al. (2007); Vinck et al. (2013); Senzai et al. (2019), but see Vigneswaran et al. (2011); Dasilva et al. (2019)). In agreement with previous findings, BW cells had lower baseline firing-rates than NW cells (mean BW FR = 2.5564, mean NW FR = 3.4101, P = 0.0153). Next, we analyzed the long-range phase-locking between V4 cell types and the V1 gamma rhythm. Surprisingly, only NW neurons showed a gamma-frequency peak in the V4-spike-to-V1-LFP phase-locking spectrum (Fig. 2B). This difference was not explained by the lower spiking rates of BW cells, because we used a metric unbiased by firing rate, and the difference was also observed when phase-locking spectra were weighted by the number of spikes of each cell (Vinck et al., 2013) (Fig. 2C). The strength of spike-field locking was independent of stimulus drive for BW cells, and only weakly correlated with stimulus drive for NW cells (fig. S2A).

**Fig. 2:**
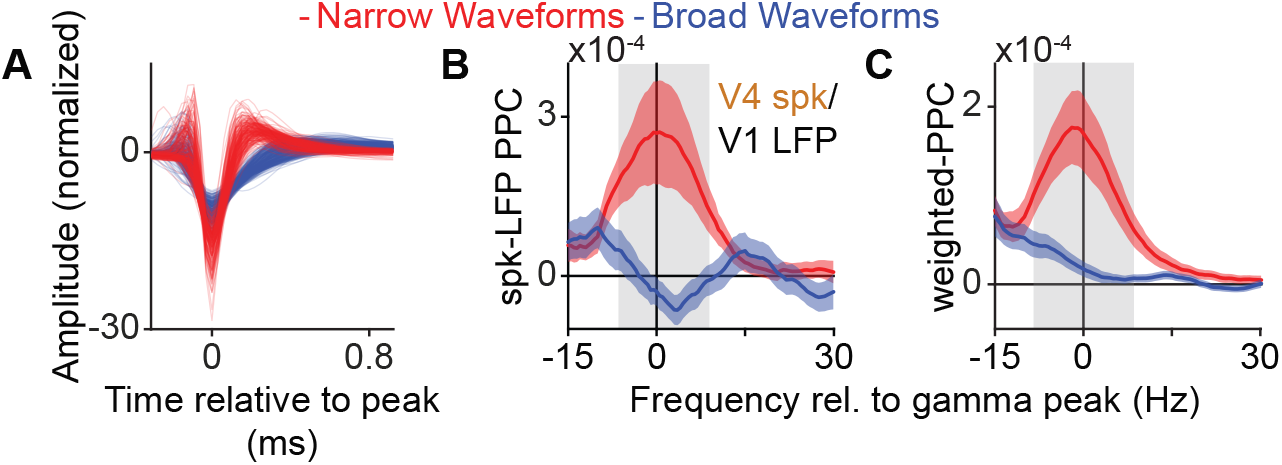
Feedforward gamma-band synchronization between V1 and V4 in the monkey mainly engages downstream interneurons. **(A)** Normalized spike waveforms of V4 SUs. **(B)** Only NW V4 neurons show gamma phase-locking to V1 LFPs (NW: *N* = 152; BW: *N* = 216). **(C)** same as B, but after weighting the PPC spectrum corresponding to each cell by the cell’s number of spikes. **(B,C)** Confidence intervals designate SEM across cells, gray rectangles designate significantly different frequency bins, randomization test between cell-types, FDR correction for multiple comparisons with a threshold of *P* < 0.05).

Together, these analyses show that long-range V1-V4 gamma-synchronization is highly cell-type specific. We wondered if the attentional modulation of V4-spike-to-V1-LFP phase-locking would therefore also be cell-type-specific. Furthermore, if the attentional modulation of V4 firing rates depends on an increase in V1-to-V4 signal transmission, then this would predict a differential attentional rate modulation of V4 NW and BW neurons. In contrast to this prediction, we found that the firing rates of both BW and NW neurons showed a comparable increase with attention (Fig. 3A to C). The increase in spiking with attention for BW neurons was observed even though the V4-spike-to-V1-LFP phase-locking spectra of BW cells did not display gamma peaks in either attentional condition (Fig. 3E, left).

**Fig. 3:**
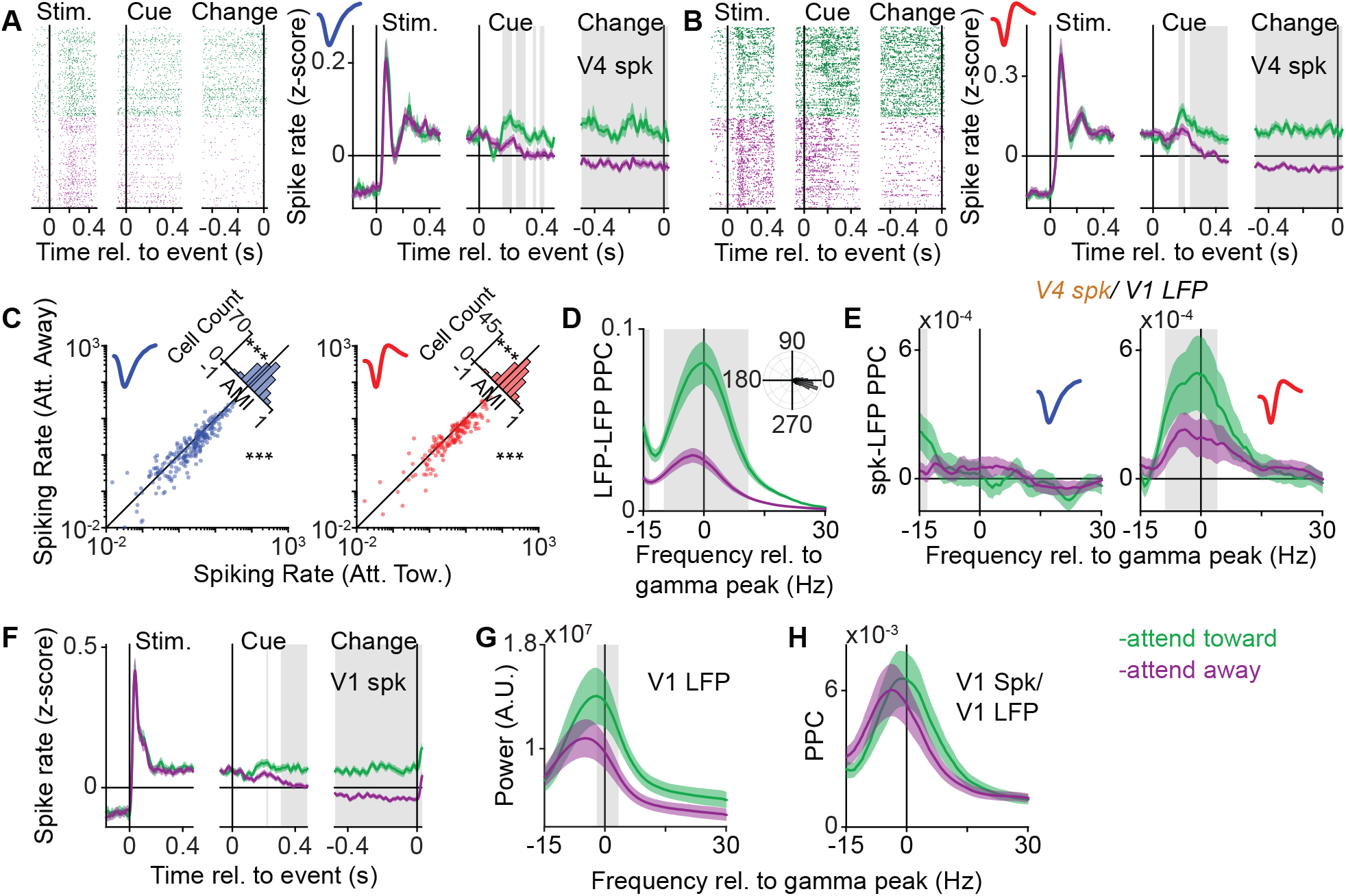
The cell-type specific effect of attention on gamma-band synchronization between V1 LFPs and V4 cells in the monkey. **(A)** Left: Example V4 BW cell. Right: Average peri-event time histogram (PETH) across V4 BW cells (**N* =* 240). **(B)** Same as A, but for V4 NW cells (*N* = 164). **(C)** Mean firing rates for attend toward and away. Insets: Attentional Modulation Index (AMI). **(D)** Phase locking (PPC) between V1 and V4 LFPs. Inset: Relative phase difference between attention conditions across all V1 and V4 LFP pairs (mean *angle* = −0.098 radians). **(E)** Phase-locking (PPC) between V1 LFPs and V4 BW (*N* = 162) and NW (*N* = 134) spiking. **(F)** Same as Fig. A (right) but for V1 SUs (*N* = 335). **(G)** Mean power of V1 LFPs for the two attentional conditions. **(H)** Same as Fig. E but for V1 LFPs and V1 SU spiking (*N* = 254). **(A,B,D-H)** Confidence intervals designate SEM across cells (A,B,E,F,H) or sessions (D,H, *N* = 68). Gray rectangles designate significantly different time or frequency bins, randomization test between cells or sessions, FDR correction for multiple comparisons with a threshold of *P* < 0.05). **(C)** \*\*\**P* < 0.001; randomization test between attentional conditions, Wilcoxon’s signed-rank test across cells.

Despite this lack of attentional modulation, there was a robust increase in gamma-band phase-locking between V1 and V4 LFPs, in accordance with previous reports (Bosman et al., 2012; Grothe et al., 2012) (Fig. 3D). Different from V4 BW neurons, we found that NW cells displayed stronger spike-to-V1-LFP gamma-band phaselocking with attention (Fig. 3E, right). This difference between NW and BW neurons was not due to a potential signal-to-noise ratio discrepancy related to differences in spiking rates in the two attention conditions (fig. S3). Furthermore, the lack of attentional modulation of V4-to-V1 phase-locking in BW neurons was also found for BW neurons with a strong attentional modulation (fig. S2B).

The increase in V4-to-V1 phase-locking with attention observed for V4 NW-neurons and V4 LFPs was likely partially due to changes in activity in V1 (Schneider et al., 2021), including an increase in firing rates (Fig.3F), gamma power and peak frequency (Fig. 3G and H). Yet, the increased propagation of the V1 gamma rhythm to V4 only affected putative FS neurons, which is likely due to the difference in filtering properties between FS and excitatory neurons (Pike et al., 2000; Vaidya and Johnston, 2013).

These analyses indicate that the changes in attentional rate modulation in V4 do, for a large part, not result from increases in V1-V4 gamma-synchronization. Furthermore, the rate modulation is found in most neurons whereas phase-locking modulation only occurs in a subset of neurons which are likely located in the granular input layer (see Fig. 5 and 7). We thus predicted that V4 firing rates may allow for more reliable decoding of the attentional state than V1-V4 phase-locking. To this end, we performed decoding analyses of the attentional state separately on spiking rates or various measures of inter-areal gamma-band synchronization. The accuracy of decoding based on single-cell spiking rates was significantly above the chance level, with both cell types displaying similar accuracy (Fig. 4A). For this reason, we pooled cell types for our decoding analyses based on population spiking rates. Decoding accuracy increased monotonically as a function of the number of simultaneously recorded neurons, without reaching an asymptote for 5 or even 10 cells (Fig. 4B). This suggests that the upper bound of information related to the attentional state, as encoded in V4 population spiking, far exceeds what can be gleaned by our findings. By contrast, decoding based on the gammaphase of spiking (Fig. 4B) or the strength of spike-LFP phase locking in the gamma range (Fig. 4C) were indistinguishable from chance. The relative gamma phases of V1 and V4 LFP pairs and the strength of their phase-locking in the gamma range were significantly more informative than chance, but still less informative than even single cell spiking (Fig. 4C). Thus, selective gamma-band synchronization contains substantially less information about the attentional state, even at the population level. Further, this finding supports the conclusion that the modulation of V4 firing rates is not a consequence of an increase in V1-V4 gamma-rhythmic synchronization.

**Fig. 4:**
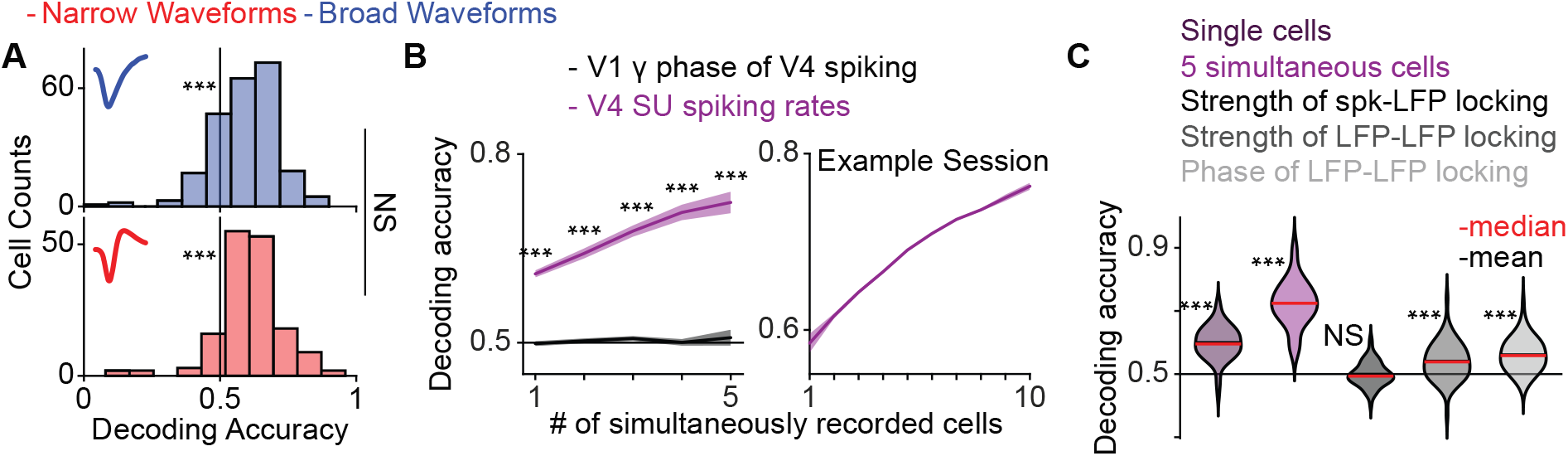
Spiking rates are more reliable in encoding the attentional state compared to measures of inter-areal gamma-band synchronization. **(A)** Decoding accuracy of the attentional condition based on the firing rates of BW (*N* = 231) and NW cells (*N* = 160). **(B)** Left: Decoding accuracy of the attention condition as a function of the number of simultaneously recorded cells for V4 SU spiking rates or the V1 gamma phase of V4 spiking, across sessions. Right: Decoding accuracy based on firing rates for an example session. **(C)** Violin plots of the distribution of decoding accuracy across sessions for decoding based on different modalities (black lines representing the mean are not clearly visible because they greatly overlap with red lines representing the median). **(A,B (left),C)** \*\*\**P* < 0.001; Wilcoxon’s signed-rank test across cells (A) and sessions (C, *N* = 68), randomization test between cell-types (A) or across sessions (B (left)). **(C)** Confidence intervals designate SEM across sessions (C (left), *N* = 68) or cells (C (right)).

**Fig. 5:**
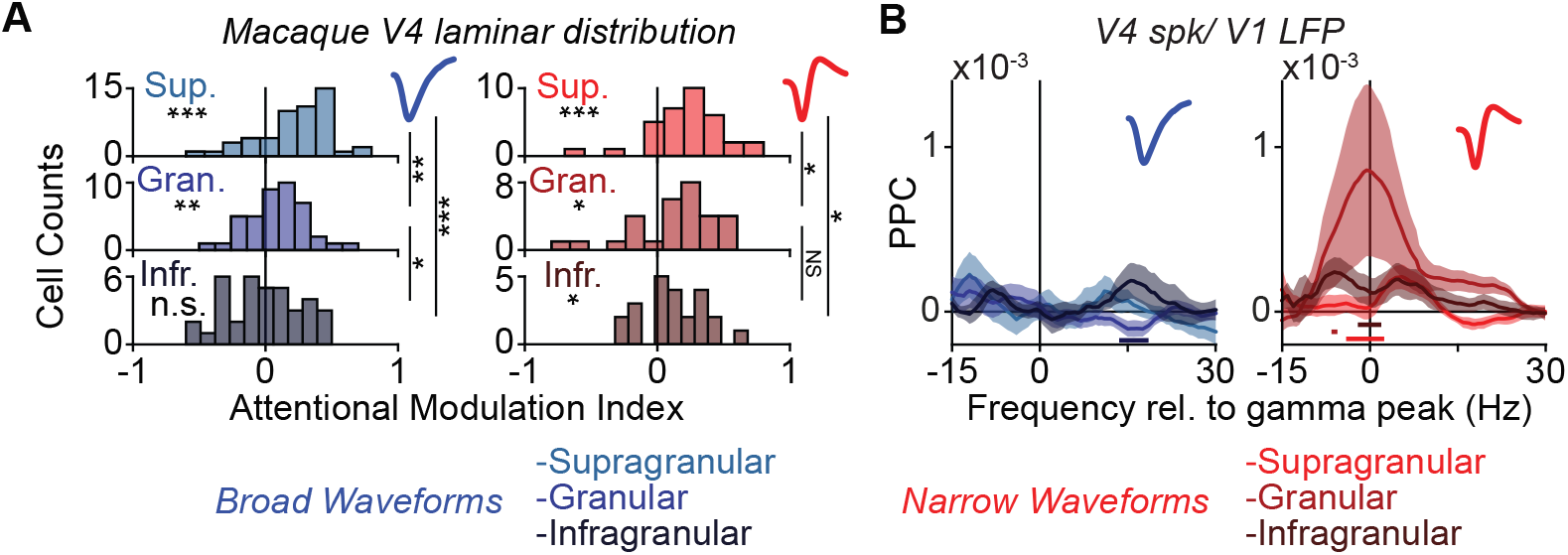
Inter-areal gamma-synchronization and attention in macaques involve cells in different laminar compartments. **(A)** AMIs at different laminar compartments for BW (left) and NW cells (right) in macaque V4. \**P* < 0.05, \*\**P* < 0.01, \*\*\**P* < 0.001; Wilcoxon’s signed-rank test across cells and randomization test between laminar compartments (BW: *Nsup*. = 52, *Ngra*. = 42, *Ninf*. = 37, NW: *Nsup*. = 34, *Ngra*. = 30, *Ninf*. = 23). **(B)** PPC between V1 LFPs and V4 cell spiking in different laminar compartments for BW (left) and NW (right) cells (BW: *Nsup*. = 43, *Ngra*. = 39, *Ninf*. = 34, NW: *Nsup*. = 30, *Ngra*. = 26, *Ninf*. = 23). **(A, B)** Bright, intermediate, and dark colors designate cells in superficial, granular, and deep layers, respectively. **(B)** Bright, intermediate, and dark colored horizontal bars in PPC spectra designate significantly different frequency bins between the superficial and granular compartment, the superficial and infragranular compartment, and the granular and infragranular compartment, respectively. Statistical comparisons are done in the same way as in Fig. 2B.

**Fig. 6:**
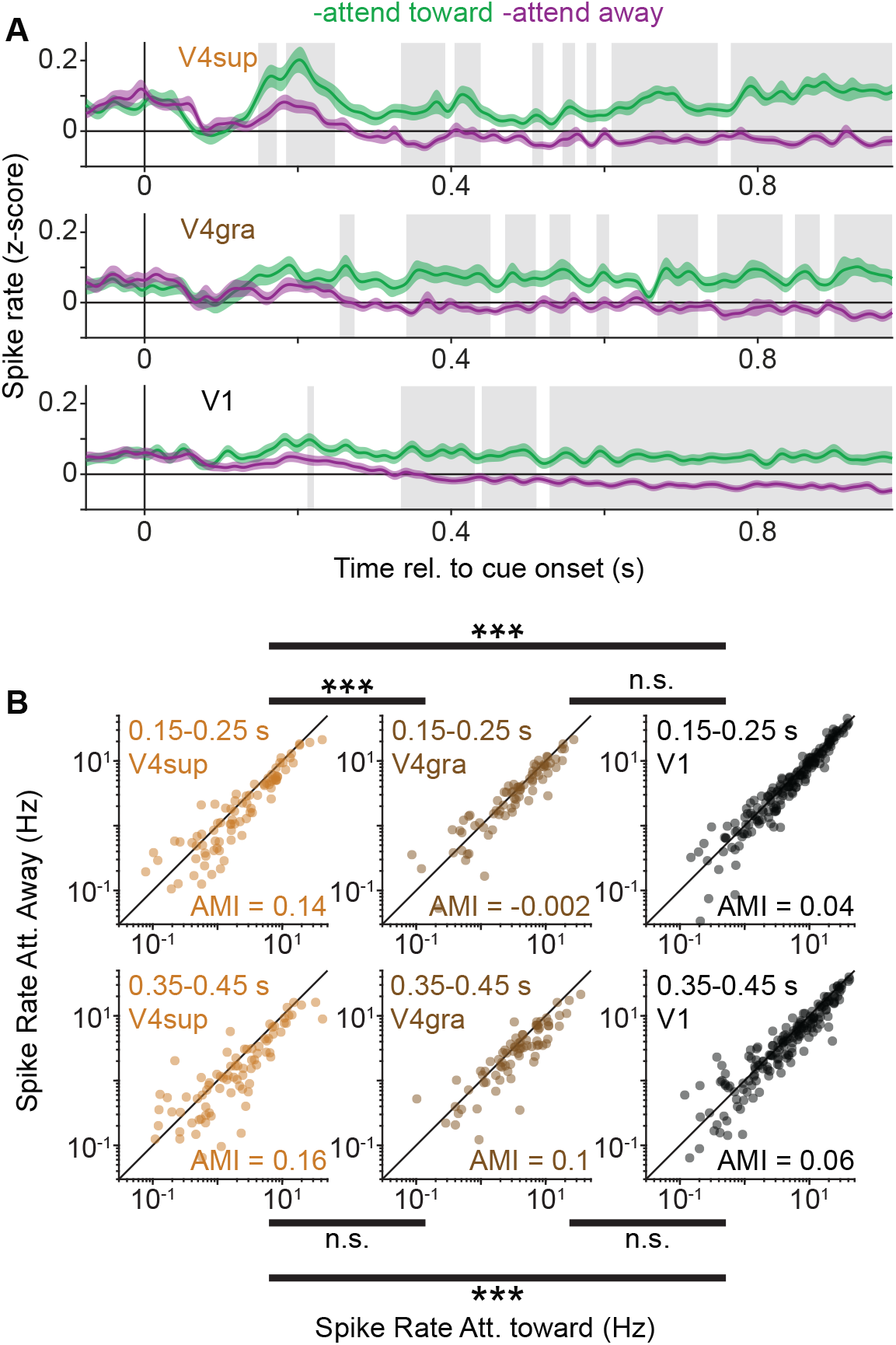
Superficial layers in V4 exhibit shorter attentional latencies in spiking compared to the input layer and V1. **(A)** Average PETH across SUs in superficial layers of V4 (top), the input layer in V4 (middle), and in V1 (bottom), for the two attentional conditions. Spiking activity is triggered around cue onset. Statistical comparisons were performed in the same manner as Fig. 2A (right). **(B)** Mean spiking rate of SUs in superficial layers of V4, the input layer in V4, and in V1 for the attend-toward (abscissa) and attend-away conditions (ordinate). Spiking rates were computed for the period between 0.15 and 0.25s after cue onset (top row) and the period between 0.35 and 0.45s after cues onset (bottom row). Randomization test between compartments; \*\*\**P* < 0.001. **(A,B)** *NV* 4*sup*. = 90, *NV*4*gra*. = 80, *NV* 1. = 181,

**Fig. 7:**
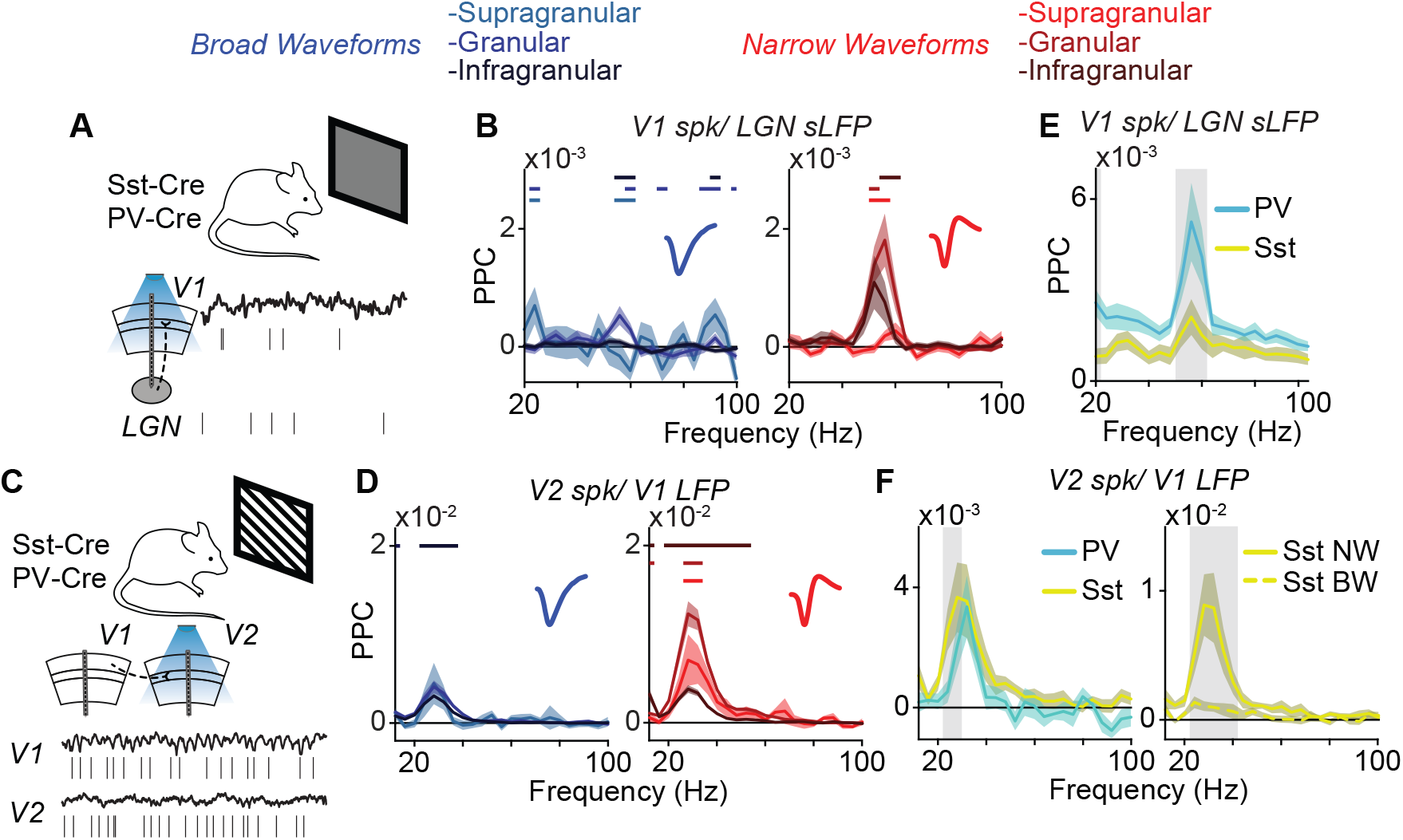
Inter-areal gamma-synchronization in mice mainly involves FS interneurons cells in feedforward input layers. **(A)** Experimental setup in mice for the luminance condition. **(B)** Same as Fig. 5A, but for V1 SUs and LGN spike-derived LFPs (sLFP) under the luminance-gamma condition, in mice (BW: *Nsup*. = 148, *Ngra*. = 463, *Ninf*. = 1055, NW: *Nsup*. = 47, *Ngra*. = 226, *Ninf*. = 316). **(C)** Experimental setup in mice for the contrast condition. **(D)** Same as (B), but for V2 SUs and V1 LFPs under the contrast-gamma condition, in mice (BW: *Nsup*. = 23, *Ngra*. = 428, *Ninf*. = 2187, NW: *Nsup*. = 11, *Ngra*. = 281, *Ninf*. = 544). **(E)** PPC between V1 LFPs and V1 cell spiking under the luminance-gamma condition, for PV+ (*N* = 47) or Sst+ cell spiking (*N* = 30). **(F)** Left: PPC between V2 LFPs and V1 cell spiking under the grating-gamma condition, for PV+ (*N* = 32) or Sst+ cells (*N* = 21). Right: PPC between V2 LFPs and V1 Sst+ cell spiking under the grating-gamma condition, for BW (*N* = 14) or NW cells (*N* = 7). **(B,D)** Bright, intermediate, and dark colors designate cells in superficial, granular, and deep layers, respectively. Bright, intermediate, and dark colored horizontal bars in PPC spectra designate significantly different frequency bins between the superficial and granular compartment, the superficial and infragranular compartment, and the granular and infragranular compartment, respectively. **(B,D-F)** Statistical comparisons are done in the same way as in Fig. 2B.

Taken together, our observations suggest that there are two distinct pathways through which V4 activity is modulated by attention: (1) an enhancement of FF V1-V4 gamma-synchronization; (2) a modulation of V4 firing rates independent of V1-V4 gamma-synchronization. We predicted that these two mechanisms should act on different laminar compartments, respectively: (1) The granular layer 4, in which FF inputs arrive, and (2) the supragranular layer, which receives prominent top-down FB (Markov et al., 2014; Vezoli et al., 2021a; Ferro et al., 2021b). To investigate this, we compared the laminar patterns of the attentional modulation of spiking and the phase-locking of V4 spiking and V1 LFPs. As predicted, we observed the strongest effects of attention in the spiking of superficial layers, for both cell types. The strength of this attentional modulation decreased sharply with depth for BW cells, whereas it was more uniform across layers for NW cells (Fig. 5A). Further, SUs in the superficial layers of V4 exhibited an earlier attention-dependent increase in their spiking compared to SUs in both the input layer of V4 and area V1 (Fig. 6). By contrast, consistent with the FF propagation of the V1 gamma-rhythm, the phase-locking of V4 cells to V1 gamma-range LFPs was strongest in the granular input layer and only present for NW cells (Fig. 5B). The laminar profile further suggests that V4-to-V1 gamma-phase locking did not propagate to the superficial layers of V4 (Fig. 5B). These findings support the existence of two separate pathways for attentional modulation, namely a FF pathway targeting L4 of V4, and a FB pathway targeting L2/3 of V4.

To physiologically characterize interneurons via optogenetic tagging and generalize these findings, we analyzed simultaneous recordings in the lateral geniculate nucleus (LGN) and areas V1 and V2 of awake mice under conditions of visual stimulation (experiments were performed by the Allen Institute for Brain Science) (Fig. 7A and C). We compared two conditions of passive visual stimulation during active states (i.e. high arousal and locomotion): (1) In the “luminance” condition, mice viewed a gray screen, which gives rise to a faster gamma rhythm (~ 60*Hz*) generated in the LGN (fig. S4A; see also (Saleem et al., 2017; Schneider et al., 2021)); (2) in the “contrast” condition, mice viewed drifting gratings, which give rise to a slower gamma rhythm (~ 30*Hz*), thought to be generated in the superficial compartment of V1 (fig. S4B and fig. S6B; see also (Veit et al., 2017)).

Our findings in mice showed a close similarity to our observations in the macaque: (1) The strength of phaselocking of spikes to the gamma generator (LGN and V1 respectively for the two conditions) was relatively weak in post-synaptic targets (V1 and V2, respectively; fig. S4A and B); (2) spiking in the respective gamma generator occurred at an earlier gamma phase compared to the spiking of its post-synaptic targets (fig. S4, C and D); (3) both the LGN and V1 gamma preferentially engaged NW cells in the downstream area (fig. S5A, for waveforms in V1 and V2), independent of brain state (fig. S5B to G); (4) phase-locking between the gamma generator (LGN and V1 respectively) and spiking in downstream cells (V1 and V2 respectively) was concentrated in FF input-layer and strongest for NW cells (Fig. 7B for luminance gamma, Fig. 7D for contrast gamma).

We investigated the differential contribution of interneuronal subtypes to inter-areal gamma synchronization via the optogenetic tagging of units corresponding to PV+ and Sst+ interneurons in mouse V1 and V2 (Pvalb-IRES-Cre and Sst-IRES-Cre lines respectively). This analysis revealed that LGN gamma in the luminance condition predominantly engaged V1 PV+-cells (Fig. 7E), the majority of which displayed an NW phenotype (fig. S7A and B). Surprisingly, both, PV+ and Sst+ cells in V2 displayed phase locking with the V1 gamma rhythm under the contrast condition. However, a closer examination of the waveform characteristics of Sst+ cells revealed that NW, and not BW, Sst+ cells were preferentially locked to V1, suggesting that they display a fast-spiking phenotype (Scala et al., 2019) (Fig. 7F). These NW Sst+ cells were localized to the granular and infragranular layers, but not the superficial layers (fig. S7C).

## Discussion

We investigated the interactions between feedforward and feedback pathways involving distinct cell types in macaque areas V1 and V4 during an attention task. Our first main finding is a concrete and highly specific functional consequence of afferent gamma rhythms in driving L4 fast-spiking interneurons in the downstream target area. Our analyses in mice revealed a similar pattern that was specific to optogenetically identified fast-spiking PV+ and narrow-waveform SSt+ interneurons, suggesting a general mechanism in cortical communication. Our second main finding is the identification of two distinct pathways through which attention modulates neural activity. Specifically, our V1-V4 analyses in macaques established a dissociation between firing rate modulation and gamma phase-locking: Phase-locking of V4 neurons to the V1 afferent gamma rhythm was specific to L4 fast-spiking interneurons in V4, whereas the attentional modulation of firing rates in V4 was expressed by both principal and inhibitory neurons and was strongest in L2/3 instead of L4. The rate modulation in superficial layers of V4 occurred at earlier latencies than in V1 and in L4 of V4. This dissociation between feedforward and feedback effects suggests distinct pathways through which attention modulates V4 activity: (1) Via direct top-down feedback to superficial layers of area V4, which amplifies the firing rate responses to the attended stimulus in both excitatory and inhibitory neurons; (2) via an amplification of feedforward gamma-synchronization between L4 fast-spiking interneurons in V4 and the V1 gamma rhythm.

### Cell-type specific inter-areal interactions

It remains a widespread assumption that the rhythmic synchronization of spiking discharges in a pre-synaptic population increases their impact on a post-synaptic target population (Bernander et al., 1991, 1994; Salinas and Sejnowski, 2001; Bruno and Sakmann, 2006). Here we tested this assumption directly and separated the impact on post-synaptic fast-spiking interneurons and excitatory neurons. Our findings demonstrate a concrete functional consequence of afferent gamma rhythms in driving L4 fast-spiking interneurons, but not excitatory neurons. The observed difference in phase-locking properties agree well with known differences in the filtering (resonance) properties of E and I cells (Pike et al., 2000; Cardin et al., 2009; Vaidya and Johnston, 2013; Moradi Chameh et al., 2021; Beaulieu-Laroche et al., 2018). Our findings also agree with observations made in rodent CA1 (Schomburg et al., 2014), wherein gamma-rhytmic input from the entorhinal cortex and CA3 mainly drive fast-spiking interneurons in the stratum lacunosum-moleculare and the stratum radiatum, respectively. Our analyses of identified GABAergic interneurons in mice further elucidated the physiological character of the fast-spiking interneurons: High-frequency (55Hz) gamma-synchronization in the LGN recruited almost exclusively V1 PV+ interneurons, mainly in L4. Low-frequency (25Hz) gamma-synchronization in V1 lead to phase-locking in V2 PV+ interneurons, but also narrow-waveform SSt+ interneurons. It is possible that this subset of SSt+ neurons corresponds to non-Martinotti cells. These neurons, located in L4 and L5, generally have action potentials with narrower widths, exhibit higher firing rates, and mostly project locally (Xu et al., 2013; **?**). Thus, the fast-spiking interneurons in V4 may comprise a mixture of PV+ and narrow-waveform SSt+ interneurons.

We conclude that feedforward gamma-synchronization primarily engages downstream inhibitory cells. In our view, this finding reflects the feedforward amplification of inhibition in the downstream area: Rhythmic input with energy in higher frequencies increases the gain of pre-synaptic spikes onto FS interneurons, due to the resonance properties of the latter (Pike et al., 2000; Cardin et al., 2009; Izhikevich et al., 2003). The lack of propagation of V1 gamma-synchronization beyond L4 of V4 contradicts the common conception that inter-areal LFP phase-locking reflects synchronization between relatively large neural populations (Fries, 2015; Palmigiano et al., 2017), considering that neurons in L4 do not make feedback projections. Furthermore, the observed lack of phaselocking of V4 excitatory neurons to V1 gamma suggests an entirely different interpretation of inter-areal phase-locking than the communication-through-coherence hypothesis, in which gamma-synchronization is thought to facilitate communication to excitatory projection neurons (Fries, 2015). Rather, the resulting increase in feedforward inhibition due to gamma-rhythmic inputs likely increases the signal-to-noise ratio of excitatory cells via a divisive operation (Atallah et al., 2012; Wilson et al., 2012; Carandini and Heeger, 2012), and may enhance the capacity of these cells to transmit their signals to their post-synaptic targets (Sohal et al., 2009; Hamilton et al., 2013). This effect may be especially pronounced for stimuli with a high degree of spatial and temporal predictability. Visual stimuli with these properties elicit strong gamma-band synchronization in macaque V1 (Peter et al., 2019a; Uran et al., 2022; Hermes et al., 2019), which may subsequently lead to a disproportionate increase of inhibition in downstream areas such as V2 and V4, thereby gating the feedforward flow of redundant information (Uran et al., 2022). A complementary role for feedforward gamma-synchronization and the concomitant feedforward inhibition has been demonstrated in the circuit comprising the entorhinal cortex, dentate gyrus and area CA3. In this circuit, inhibitory feedback was shown to be important for pattern separation (Guzman et al., 2021). It is possible that similar computations are facilitated in visual cortex via the modulation of inter-areal gamma synchronization by processes such as attention, which requires also the separation between multiple competing inputs (Desimone et al., 1995).

### Distinct feedforward and feedback pathways for attentional modulation

Our results demonstrate a dissociation of feedforward and feedback influences in V4 under attention. We show that feedforward gamma-rhythmic influences from V1 do not propagate further than the V4 input layer, and they primarily recruit FS interneurons, which principally make projections to local neurons. By contrast, we observed attentional increases in firing rate for, both, excitatory cells and FS interneurons in all layers of V4. Crucially, these attention-related increases appear earliest in the superficial layers of V4, and are, therefore, not brought about by feedforward input, but rather by top-down feedback (Douglas and Martin, 2004; Chaudhuri et al., 2015; Vezoli et al., 2021b; Felleman and C Van Essen, 1991). Feedback projections are likely modulatory, that is they do not induce spiking, rather they amplify the gain of their post-synaptic targets. The mechanism of this amplification remains to be determined: One possible hypothesis is that top-down modulation is mediated by disinhibition, i.e. the inhibition of Sst+ interneurons following the long-range activation of VIP+ interneurons (Shen et al., 2022; Batista-Brito et al., 2018; Zhang et al., 2014; Pfeffer et al., 2013). As SSt+ neurons project to both PV+ and excitatory cells (Gentet et al., 2012), attention would lead to an increase in the gain of both fast-spiking interneurons and excitatory cells, as observed in this study. Likewise, it remains to be investigated what mechanism accounts for increased phaselocking in V4 fast-spiking interneurons to the V1 gamma rhythm. One possibility is that attention enhances the excitability of V4 fast-spiking interneurons, e.g. through the inactivation of SSt+ interneurons. Another possibility is that the increase in V1 gamma power and firing rates lead to increased phase-locking, such that the latter reflects enhanced input power at the V1 gamma-frequency (Schneider et al., 2021).

The existence of two pathways raises the question of their relative importance. To investigate this, we performed population decoding analyses, and established that V4 firing rates are substantially more reliable than the phase-locking strength or the relative phase of V1 and V4 LFPs in encoding the locus of attention. This is the case even at the level of single cells, with a notable increase in reliability when considering multiple simultaneously recorded neurons. It is instructive to consider how a noisy signal (such as the firing rate of a single neuron) can be more reliable than a population-wide signal (such as LFPs in V1 and V4) in encoding a population-wide modulatory state such as attention. A modeling study by Akam and Kullmann (2012) has shown that a modulation of coherence (increase for attended target) is unable to reliably achieve selective communication with a downstream receiving population. This is the case because input from a non-selected sender population will, by chance, often arrive at downstream phases of high excitability, even if the selected sender is perfectly phase-locked with the receiver Akam and Kullmann (2012). Altogether, this analysis supports the idea that the firing rate modulation in superficial layers mediated by feedback, instead of the increase in feedforward V1-V4 gamma coherence, is the primary attentional selection mechanism.

### Conclusion

These findings provide, to our knowledge, the first description of inter-areal interactions between distinct cell types residing in separate cortical layers, while animals perform a cognitive (attention) task. Our analyses revealed a motif that we also confirmed in mice, namely the recruitment of fast-spiking interneurons in a downstream target area via gamma-synchronization. Future work should identify the precise biophysical mechanisms underlying the phenomenon, and further investigate the generality of this observation. Furthermore, our data show that inter-areal gamma phase-locking and rate modulation may constitute distinct mechanisms for top-down attention. Altogether, these findings reveal distinct, cell-type-specific feedforward and feedback pathways for the attentional modulation of inter-areal synchronization and spike rates, respectively.

## Acknowledgments

This study was supported by an ERC Starting Grant [SPATEMP, EU] (M.V.), a BMBF (Germany) Grant [Computational Life Sciences, project BINDA, 031L0167] (M.V.), a grant from the Wellcome Trust [093104] (J.v.K., A.T.), and two grants from the MRC: [MR/P013031/1] (J.v.K., M.A.G., A.T.), and [MR/K013785/1] (M.A.G., A.T.).

## Methods

### Animals and experimental procedures: Macaques

Our study involved two male adult rhesus macaque monkeys (Macaca mulatta, age 10-12 years, weight 8.5-12.5 kg), implanted with a head post and recording chambers over areas V1 and V4 under sterile conditions and general anesthesia. Housing conditions, surgical procedures, and post-operative care conditions have been described in considerable detail in previous papers (Gray et al., 2016; Thiele et al., 2006) and were in accordance with the UK Animals Scientific Procedures Act, the National Institute of Health’s Guidelines for the Care and Use of Animals for Experimental Procedures, and the European Communities Council Directive RL 2010/63/EC.

### Recordings in Mice

Analyses in mice were based on the publicly available Visual Coding - Neuropixels dataset, which was collected and preprocessed by the Allen Institute for Brain Science (Siegle et al., 2021). This electrophysiology dataset comprises single unit spiking and LFP signals recorded simultaneously from 4 to 6 visual areas in 57 awake mice, under conditions of visual stimulation by various stimuli. In this study we focused on the interactions of the lateral geniculate nucleus (LGN), area V1, and area V2 (comprising the lateral (VISl), anterolateral (VISal), rostrolateral (VISrl), anteromedial (VISam), and posteromedial (VISpm) visual areas). Surgical procedures, visual stimulation protocols, recording equipment/techniques, signal preprocessing, and spike sorting have been extensively described in the technical white paper accompanying the dataset (https://portal.brain-map.org/explore/circuits/visual-coding-neuropixels), and will not be discussed here.

### Behavioral paradigm: Macaques

Stimulus presentation and behavioral control were performed by using the Remote Cortex 5.95 software (Laboratory of Neuropsychology, National Institute for Mental Health, Bethesda, MD). We presented visual stimuli on a cathode ray tube (CRT) monitor, with a refresh rate of 120 Hz, a resolution of 1280 × 1024 pixels, at a distance of 54 cm from the eyes of the macaques, under conditions of head-fixation. The macaques performed a standard selective visual attention task described in more detail in (van Kempen et al., 2021). In brief, each trial was initiated when the macaques held a lever and foviated a white fixation spot (0.1°) displayed at the center of the screen on a gray background (1.41 cd/m2). Releasing the lever or breaking fixation for the duration of the trial led to the trial’s termination. After a pre-stimulus fixed delay period (duration of 614 and 674 ms, for macaques T and W, respectively), three peripheral colored square-wave gratings were presented at the same eccentricity and the same distance from each other, with one of the stimuli being centered on the RFs of neuronal populations in the V1 and V4 recording sites. We adjusted the diameter of the presented stimuli based on eccentricity and size of RFs, with stimulus diameters ranging from 2 to 4°. The color of each grating was pseudo-randomly permuted between recording sessions but remained fixed for the duration of each session. The gratings in the majority of recordings drifted perpendicular to the orientation of the grating, with the motion direction pseudo-randomly assigned on every trial. In 22 out of the total of 34 sessions, macaque W was shown stationary gratings. After a random delay (618-1131 ms for monkey T, 618-948 ms for monkey W; period duration chosen from a uniform distribution), a central colored cue appeared, matching the color of one of the peripheral gratings, thereby designating the target stimulus for the trial. The cue color was randomly assigned for each trial. At a random delay after the presentation of the cue, the luminance of one of the peripheral stimuli decreased (1162-2133 ms for macaque T, 1162-1822 ms for macaque W; period duration chosen from a uniform distribution). If the luminance change occurred on the target stimulus, the macaque was rewarded for subsequently releasing the lever. If, however, the luminance change occurred on one of the other two stimuli, the macaque was only rewarded after maintaining fixation and keeping hold of the lever until a luminance change occurred on the target stimulus, which corresponded to either the second or third luminance change event (each event following the previous luminance change after 792-1331 ms for monkey T and 792-1164 ms for monkeys W; period duration chosen from a uniform distribution).

### Data acquisition: Macaques

Daily electrophysiological recordings from all cortical layers of visual areas V1 and V4 were performed using 16-contact linear electrode shanks (150 micrometer contact spacing, Atlas silicon probes), by inserting the shanks perpendicular to the cortex. Raw data were collected using an HS-36 Neuralynx pre-amplifier and a Neuralynx Digital Lynx amplifier with the Cheetah 5.6.3 data acquisition software interlinked with Remote Cortex 5.95. Data were sampled at 24 bit with a 32.7 kHz sampling rate and stored to a disc. We recorded eye position and pupil diameter at a rate of 220 Hz with a ViewPoint EyeTracker (Arrington Research).

### Analysis Software

The analyses presented in this study were performed in MATLAB (The MathWorks) and used custom scripts and the FieldTrip analysis toolbox (https://www.fieldtriptoolbox.org).

### Receptive Field Mapping: Macaques

The estimation of electrode receptive fields (RFs) was based on the envelope of MUA (MUAe). This signal was extracted after low-pass filtering (5th order Butterworth filter with a corner frequency of 300Hz) the rectified 0.6-9 kHz filtered signal. RF mapping involved the presentation of 0.5–2° black squares on a 9 × 12 grid. An offline response map was computed for each channel via the reverse correlation of the MUAe signal to these stimuli. Next, this map was then converted to z-scores, and RFs for each channel were defined as the region surrounding the peak activity that exceeded a z-score of 3. More detailed information about our approach in RF estimation can be found in (Gieselmann and Thiele, 2008).

### Preprocessing: Macaques

In macaques, we extracted local field potentials (LFPs) from the broadband signal by low-pass filtering (6th order Butterworth filter with a corner frequency of 500 Hz), high-pass filtering (3rd order Butterworth filter with a corner frequency of 2 Hz) and down-sampling to ~ 1.0173Hz. A Butterworth bandstop filter (50 Hz and harmonics ±0.2 Hz) was additionally used to remove powerline artifacts at 50 Hz and harmonics. Spike waveforms were extracted from the broadband signal after taking the following steps: 1) The median-filtered signal (3ms window) was subtracted from the broadband signal. 2) This high-pass filtered signal was further band-pass filtered (second order Butterworth filter with corner frequencies of 20Hz and 8kHz). 3) The signal was further de-noised by subtracting the shank-wide median signal from each electrode in the corresponding electrode shank. 4) Individual spikes were detected as negative voltage crossings of a threshold of 5 absolute deviations. 5) Windows surrounding the local minima of these negative crossings were examined for the presence of early large positive peaks or double negative peaks (window length of ±20 samples), and spikes with these characteristics were discarded. 6) Individual spike waveforms were defined as data points spanning −15 to 30 samples around the local minima that were not discarded.

### Spike Sorting: Macaques

We isolated single units (SUs) from macaque visual cortex in a semi-automated manner. Semi-automatic clustering was performed with the KlustaKwik software (version 1.7) on the following features: 1) The energy of the spike waveform and the energy of its first derivative. 2) The four principal components and the three Haar wavelet components explaining most of the variance of all detected spike waveforms. MClust (version 3.5) was used to manually assess the quality of isolation of each candidate cluster. Clusters were deemed to correspond to an isolated single unit if the following criteria were met: 1) The isolation distance (ID) of the cluster from the noise cluster (Schmitzer-Torbert et al., 2005) exceeded a value of 15. 2) The L-Ratio (LR) (Schmitzer-Torbert et al., 2005) of the cluster was lower than 0.2. 3) Less than 0.05% of interspike intervals were below 1 ms. The population of single units that were included in our analysis had a median ID of 26.1013 and median LR of 0.0406.

### Waveform Classification

The mean waveforms used in waveform classification were extracted by applying a median filter (window length of 3ms) on the broadband voltage trace, subtracting the median-filtered signal from the raw broadband signal, and computing the average across data segments of −20 to 96 samples around the timepoint corresponding to each SU spike. The mean waveform of each SU was then normalized by subtracting the median of the first and last 10 samples in the waveform, and subsequently dividing by the waveform’s energy. Triphasic waveforms characterized by a stronger early rather than late positive peak, and waveforms with a DC difference between their beginning and end segment were discarded. Two-dimensional t-Stochastic Neighbor Embedding (t-SNE; perplexity of 30) was then applied on the remaining waveforms. Lastly, fuzzy c-means clustering (fuzzifier of 2, euclidean distance metric, tolerance of 10^-10^) was performed on the t-SNE matrix, which resulted in two clearly separate waveform clusters, corresponding respectively to broad and narrow SU waveforms.

### Spectral Analysis

Analyses involving LFPs, such as the computation of spectral power, and inter-areal LFP-LFP phase-locking, may be affected by the presence of electrode-headstage-related noise and the influence of the electrode reference (Pesaran et al., 2018). In this study, we addressed this issue by subtracting the average LFP signal across each recording electrode shank from the LFP signal of each corresponding electrode, in a trial-by-trial manner. After this preprocessing step, LFP-exclusive analyses were based on the multiplication of LFP data epochs of 0.5 s with seven distinct prolate Slepian tapers, and the subsequent application of the fast Fourier transform (FFT) (Mitra and Pesaran, 1999). Spectral power was computed by rectifying the resulting complex Fourier coefficients and raising them to the power of 2. The power spectra shown in Fig. 3G were produced by multiplying power values, corresponding to different frequency bins, to the square of their respective frequencies. The phase-locking strength between LFPs in V1 and V4 was quantified with the Pairwise Phase Consistency (PPC) metric, which avoids pitfalls associated with standard measures such as the number of LFP epochs corresponding to each condition (Vinck et al., 2012). In brief, PPC is computed in the following steps: 1) Phases of the V1 and V4 LFP signal are extracted from the complex Fourier coefficients corresponding to different trial epochs, and the V1-V4 relative phase is computed for each epoch. 2) The resulting complex vectors are normalized by their amplitude to produce unit vectors. 3) Conjugate multiplication is performed for each possible vector pair, and this product is then averaged and rectified. Note, that the peak frequency of stimulus-induced gamma depends on stimulus properties (Roberts et al., 2013; Ray and Maunsell, 2010; Peter et al., 2019b) (fig. S1A) and individual subjects (van Pelt et al., 2012) (fig. S1B). For this reason, we aligned mean power and V1-V4 LFP-LFP PPC spectra to the individual gamma peak of LFP-LFP phase locking separately for each session.

### Filtering for gamma-band phase extraction and trial-by-trial estimation of phase-locking strength

In macaques, the estimation of the gamma-band LFP phases of spiking (Fig. 1G), trial-by-trial spike-LFP phase-locking strength, trial-by-trial interareal LFP phaselocking strength, and the relative gamma band phases of V1 and V4 LFPs (the latter four used in decoding analyses; Fig. 4B and C) was based on using a two-way, bandpass Butterworth filter with an order of 12. As noted above, the two macaques used in this study exhibited different gamma-band peak frequencies, thus we used different corner frequencies for each macaque (40-90 for monkey T, 25-55 for monkey W). Next, we performed the Hilbert transform on the filtered data. Note that filtering and the Hilbert transform were applied on the complete trial before selecting relevant trial epochs for further analysis. The results shown in Fig. 1G were produced by collecting the time points of the occurrence of each SU spike and the computation of the mean angle across all concurrent V1 Hilbert-transformed LFP datapoints. See the ‘Decoding Analyses’ section below for further information about the analyses of filtered signals in decoding analyses.

In mice, LFP phases of SU spiking were extracted from the complex Fourier coefficients corresponding to the gamma-band peak frequency in the SU-LFP PPC spectrum.

### Quantification of Spike-LFP Phase-Locking: Macaques

Spike-LFP phase locking in macaques was assessed for the period between post-stimulus onset and target change for Fig. 1D to F, Fig. 2B and C, and Fig. 5B (period between cue onset and target change for Fig. 3E and H, and fig. S3). In these analyses, we used bipolar derivatives of the LFP, in order to mitigate the effect of 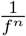 noise and lower frequency oscillations on the estimation of gamma-band spike-LFP phase-locking strength. The strength of spike-LFP phase locking was assessed with PPC, and in particular the PPC1 measure (Vinck et al., 2012), which removes bias associated with differences in spiking rates. In short, the LFP-phase of SU spiking for each frequency f, was ascertained by collecting LFP data segments of a duration of 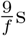 centered around each spike. We, then, multiplied these LFP segments with a Hann taper of a corresponding length, and performed an FFT on the resulting product. Only SUs with >= 200 spikes per condition were included in our analyses. This resulted in the inclusion of a different number of cells in different analyses, e.g. spectra corresponding to NW cells in Fig. 2B and Fig. 3E involved different numbers of cells. We performed additional analyses in order to control for the effect of spiking rate on spike-LFP phase locking, when comparing different cell types and attention conditions (fig. 2C and fig. S3, respectively). For the comparison between the spike-LFP phase-locking strength of BW and NW cells, we separately weighted the PPC spectrum of each cell by the ratio of its spike count and the mean spike count of each cell in the same cell class. For the comparison between attention conditions, the PPC spectrum of each cell per condition was weighted by the ratio of its spike count for the respective condition and the mean total spike count of each cell included in the analysis.

### Quantification of Spike-LFP Phase-Locking: Mice

In mice, spike-LFP phase locking for the luminance condition was assessed as in Schneider et al. (2021). Briefly, data periods during the presentation of a gray screen were divided into 1s pseudo-trials. Phase locking in the contrast condition was estimated for the period between the onset and offset of each grating stimulus. Note, that the LGN lacks the ordered columnar structure found in cortical areas and, thus, does not produce an LFP signal. Therefore, in order to examine population-wide oscillatory activity in the LGN, we estimated a surrogate LFP (sLFP) signal derived from population spiking (Schneider et al., 2021). This was done by summing the spikes of all individual isolated units in the LGN, and filtering this signal between 1 and 100Hz. We exclusively analyzed sessions where at least 10 single units were recorded in LGN. As in the macaque, spike-LFP phase locking was quantified by the PPC1 metric. This was done after segmenting the LFP in data epochs of 250ms, multiplying these segments with a Hann taper and performing the Fourier transform.

### Drivenness, attentional effect in spiking, and stimulus selectivity indices

SU peri-event time histograms (PETHs) for different trial epochs (Fig. 3A, B and F, and Fig. 6A) were computed by taking the following steps: 1) We counted the total number of spikes in bins of 1ms for the full trial. 2) The resulting time series were convolved with a gaussian kernel with a length of 50ms, and a standard deviation of 8ms. 3) We computed trial-wise z-scores of the smoothed time series. 4) Relevant trial epochs were collected and averaged across trials.

SU drivenness by the stimulus (fig. S2A) was estimated by using the following equation:

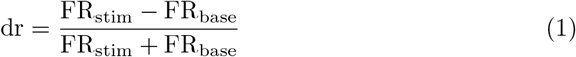

where *FR_stim_* and *FR_base_* respectively designate the mean spike-count in the period between 0.05-0.25s after stimulus onset and a period spanning 0.2s before stimulus onset. The attentional modulation index (AMI) was computed based on the following equation:

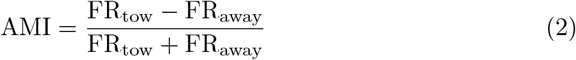

where *FR_tow_* and *FR_away_* designate the mean spikecount in the 1s period before the first stimulus change across trials (Fig. 3C, Fig. 5A, and fig. S2B; 0.15s-0.25s and 0.35-0.45s after cue onset in Fig. 6B), respectively for the attend-toward and attend-away conditions. The spiking rates shown in Fig. 3C and used in decoding analyses were computed for the same time period. Indices quantifying the selectivity of SUs, recorded in mice, to grating properties (orientation, direction, and temporal frequency) were included in the Visual Coding - Neuropixels dataset provided by the Allen Institute for Brain Science.

### Decoding Analyses: Macaques

Decoding of the macaque’s attentional state was focused on the time period spanning 1s before the onset of the first luminance change event. In our decoding analyses, we compared the decoding accuracy of 5 physiological measures: 1) SU spiking rates. 2) The strength of V1-V4 gamma-band LFP-LFP phase-locking. 3) The relative phase of V1 and V4 gamma-band LFPs. 4) The strength of the gamma-band phase-locking of V4 SU spiking to V1 LFPs. 5) The phase of V1 gamma-band LFPs in which V4 SU spiking occurred. Note that decoding analyses involving LFPs were performed on filtered and Hilbert-transformed signals, as described in section ‘Filtering for gamma-band phase extraction and trial-by-trial estimation of phase-locking strength’. The strength of phase-locking between LFPs in each trial was estimated by computing the Phase Locking Value (PLV) between the gamma-band complex time series corresponding to each 1s epoch for all possible V1-V4 LFP pairs. The trial-by-trial relative phase between each V1 and V4 LFP pair was computed after performing a timepoint-by-timepoint conjugate multiplication between the two complex vectors corresponding to each LFP pair, averaging the values in the resulting complex vector, and estimating its angle. The strength of phase-locking between V4 spiking and each gamma-band LFP in V1 was assessed in each trial by collecting the complex coefficients of the LFP corresponding to each V4 spike in the trial and computing PPC between them. Spikes in this analysis were pooled across simultaneously recorded SUs. Lastly, the V1 gamma-band phase of V4 SU spiking was estimated in each trial by computing the mean angle of complex LFP coefficients corresponding to each V4 SU spike for every SU-LFP pair.

Our decoding analyses used a maximum-likelihood (ML) estimation algorithm described in more detail in Montijn et al. (2014). In short, the ML algorithm depends on Bayes’ rule:

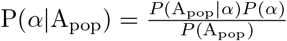

where *P*(*α*), is the prior, which designates the prior probability of the trial corresponding to the attend-toward condition; *P*(*α*|A_pop_) is the posterior, which designates the posterior probability of the trial’s neural activity A_pop_ being observed because the trial belongs to the attend-toward or the attend-away condition; *P*(A_pop_|*α*), is the likelihood, which designates the probability that the attentional condition will result in neural activity A_pop_; and *P*(A_pop_) is the model evidence, which is the probability of observing neural activity pattern Apop. For our analyses, we used only flat priors.

As mentioned above, we performed decoding analyses by either using angular quantities (relative phase of V1 and V4 LFPs or the LFP phase of spiking), the spiking rate, or other non-angular quantities (strength of phaselocking between V1 and V4 LFPs, or strength of phaselocking between spiking and LFPs). We analyzed angular quantities for any neural signal i by approximating the likelihood distribution for the allocation of attention to either the stimulus driving V1 and V4 neural activity or one of the other two stimuli by a von Mises distribution with a location *μ* and concentration *κ*. For spiking rates and non-angular quantities the likelihood was approximated, respectively, by a Poisson distribution with an expected occurrence rate λ and a Gaussian distribution with a mean *μ* and standard deviation *σ*. The decoder was trained in a leave-one-trial-out jackknife fashion.

For decoding based on population activity, we focused on the spiking rates of simultaneously recorded SUs, all possible V1 and V4 LFP pairs, and the phases of spiking of simultaneously recorded SUs in V1 gamma-band LFPs. Decoding accuracy was estimated as a function of the number of simultaneously recorded SUs by random subsampling as in Montijn et al. (2014). The posterior probability distribution for the attend-toward condition corresponding to a population of n signals can be estimated by computing the product over the posterior probabilities for all n signals:

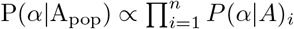

The decoded attentional condition is then defined as the condition with the largest population-wide posterior probability.

### Assignment of cortical layers

The assignment of neuronal activity to either the superficial, granular or deep laminar compartment was primarily based on the extraction of the current source density (CSD) signal from the stimulus-evoked LFP signal. In mice, CSD was extracted under conditions of whole screen flash stimulation, whereas in macaques this was done during the period after the onset of the peripheral grating stimuli used in the attention task. Stimulus-evoked CSD was computed by taking the second discrete spatial derivative of the LFP across different electrodes in each electrode shank (Mitzdorf, 1985). LFPs from electrode contacts with a relatively low signal-to-noise ratio were discarded and exchanged with a signal derived from a linear interpolation between the contacts’ neighboring electrodes. Further, in macaques, recording sessions that displayed a substantial drift of RF location with cortical depth were excluded from our laminar analyses.

In addition to the identification of sinks and sources in the stimulus-evoked CSD, different cortical laminar compartments were identified in the macaque through the inspection of the latency of stimulus-evoked multi-unit activity at different cortical depths. Peri-stimulus time histograms in this analysis included all detected negative spikes from each electrode contact and were computed similarly to other PETHs shown in our study, but with a gaussian kernel with a length of 25 ms and a standard deviation of 4 ms. Finally, we computed the logarithm of the ratio between the total number of positive spikes and the total number of negative spikes, detected in each electrode contact. Contacts with a log-ratio close to zero were deemed to lie outside of the brain, whereas contacts with negative log-ratio were assigned to the gray matter, and contacts with a positive log-ratio were presumably located in the white matter.

### Testing for optogenetic response: Mice

Optogenetic tagging experiments were performed on mutant mice expressing Pvalb-IRES-Cre and Sst-IRES-Cre. The optogenetic stimulation consisted of 10 ms pulses (for more details, see https://portal.brain-map.org/explore/circuits/visual-coding-neuropixels). Cre-expressing cells were identified using the ZETA-test Montijn et al. (2021), a recently developed parameter-free statistical test to determine whether neurons exhibit a time-locked modulation of firing rates by a specific event. First, the ZETA test was used to test which neurons showed significantly modulated spiking activity (*P* < 0.05) within a 0.5-second window following the onset of optogenetic stimulation. Next we calculated the instantaneous peak- and trough-latencies of all significantly modulated cells. Cells were classified as optogenetically tagged if their peak latencies occurred within the 10 ms of optogenetic stimulation. To avoid misclassification due to laser artifacts, neurons with a peak earlier than 1 ms after the onset of the optogenetic pulse were discarded.

### State detection: Mice

The strength of contrast gamma in the mouse was positively modulated by arousal and locomotion, respectively (fig. S5B to G, fig. S6), therefore this analysis was focused on periods of high locomotion, in order to optimize the signal-to-noise ratio of gamma LFPs. We assessed the arousal level and running speed of mice by examining, respectively, the pupil diameter signal and speed signal accompanying the Visual Coding - Neuropixels dataset, used in our study. For the luminance condition, we detected periods of high and low arousal by normalizing the pupil diameter signal by its maximum value, separately for each recording session, and classified periods in which this signal had values between 0.65 and 0.95 as periods of high arousal (values between 0.3 and 0.55 corresponded to periods of putative low arousal). For the contrast condition, states of high and low locomotion were defined as periods in which the mouse had a running speed > 5*cm/s* and < 1*cm/s*, respectively.

### Statistical Testing

Unless otherwise stated, statistical comparisons in the study were non-parametric, two-sided, and based on 1000 randomizations (Nichols and Holmes, 2002). Randomization between means of quantities (spectra, spiking rate, decoding accuracy, AMI) measured across different cell populations/types (Fig. 1D and E, Fig. 2B and C, Fig. 4A, Fig. 5A and B, Fig. 6B, Fig. 7B, D to F, fig. S4A and B, fig. S5B to G, fig. S6, fig. S7) and trial-epochs (Fig. 1C) involved randomly exchanging the quantities under comparison between populations or epochs, while keeping the original number of values per population/epoch constant. In the case of spectra, statistical significance was achieved for frequency bins where observed differences between the mean spectra of each population were larger or smaller than the 97.5th percentile of the maximal values or the 2.5th percentile of the minimal values, respectively, across all randomized difference-spectra. This approach corrects for the false discovery rate associated with multiple comparisons. In the case of singular values per cell (e.g. spiking rate) we computed p-values by taking the following steps: 1) We computed the ratio of the mean difference between populations and the standard deviation across all randomized differences. 2) We rectified this ratio and computed its cumulative density function (CDF) value. 3) We subtracted this value from 1 and divided the difference by 2. The statistical assessment of correlations (fig. S2) involved randomly shuffling the order of the PPC values corresponding to each cell and computing Spearman’s rho 1000 times. Here, p-values were computed in a similar manner as described above, with the main difference being that we compared the empirical mean correlation to a distribution of randomized correlations. Statistical comparisons between attention conditions (Fig. 3A, B, D to H, Fig. 6A, and fig. S3 were also done similarly to what was described above, with the only difference being that randomizations were based on the random switching of at-tentional condition labels.

---

**Fig. S1:**
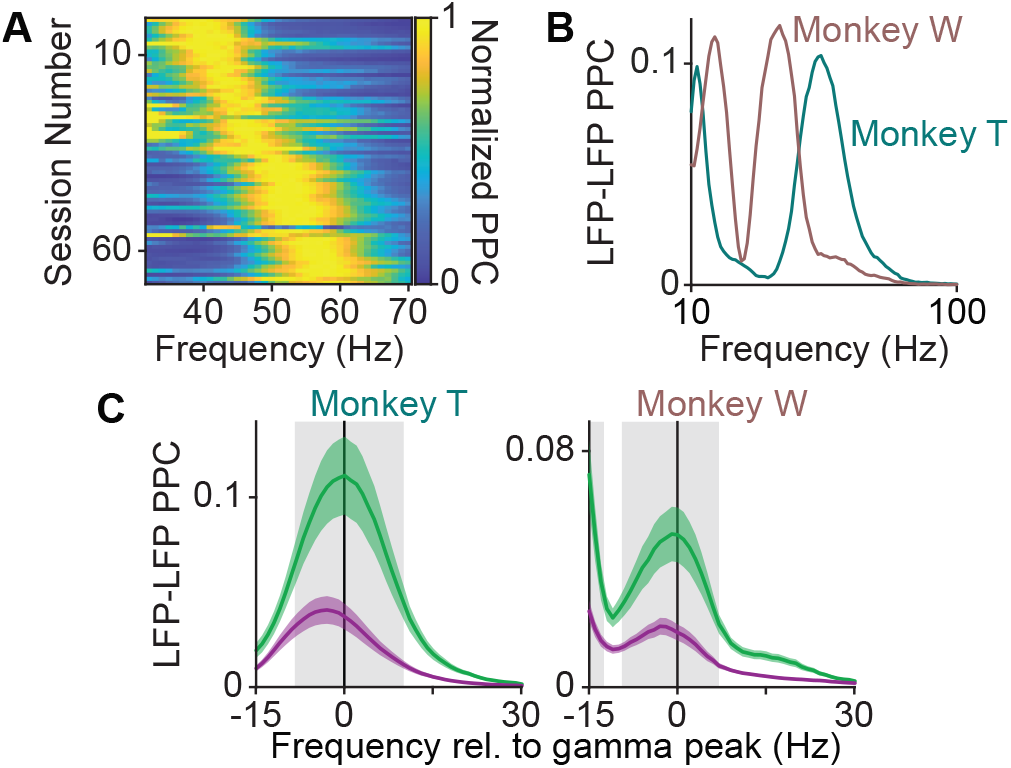
Variability in the gamma-range peak-frequency of LFP-LFP phase-locking in macaques. **(A)** Normalized LFP-LFP PPC between V1 and V4 across all inter-areal channel pairs, for all sessions (*N* = 68), sorted for gamma-range peak-frequency. **(B)** Example sessions demonstrating the difference in gamma-range peak-frequency in the two monkeys. **(C)** Same as Fig. 2D but plotted separately for the two monkeys (*N* = 34 sessions for each monkey).

**Fig. S2:**
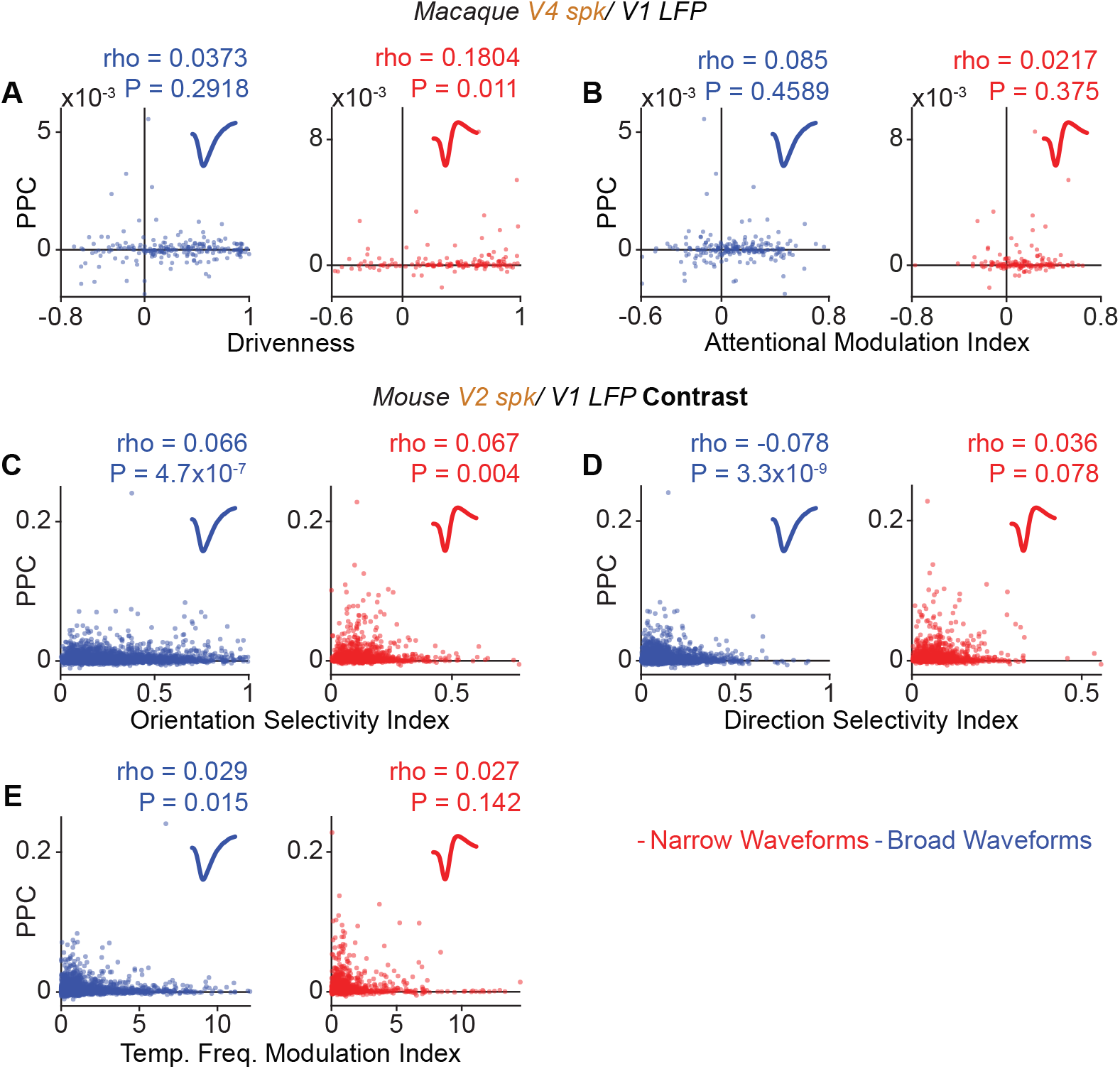
Effects of cell-drivenness and the attentional modulation of spiking on the gamma-rhythmic phase locking between downstream spiking and upstream LFPs for different cell types in macaques and mice. **(A)** Relationship between cell-drivenness and gamma-band PPC of spiking to V1 LFPs for V4 BW and V4 NW cells. **(B)** Same as A, but for the AMI of V4 spiking and gamma-band PPC of spiking to V1 LFPs. **(A,B)** BW: *N* = 216. NW: *N* = 152. **(C)** Same as A, but for the orientation selectivity index of spiking and gamma-band PPC of spiking to V1 LFPs for V2 BW and V2 NW cells, in the mouse. **(D)** Same as C, but for the direction selectivity index of spiking and gamma-band PPC of V2 spiking to V1 LFPs. **(E)** Same as C, but for the depth of modulation of spiking by the temporal frequency of the grating stimuli and gamma-band PPC of V2 spiking. **(C-E)** BW: *N* = 5477. NW: *N* = 1585.

**Fig. S3:**
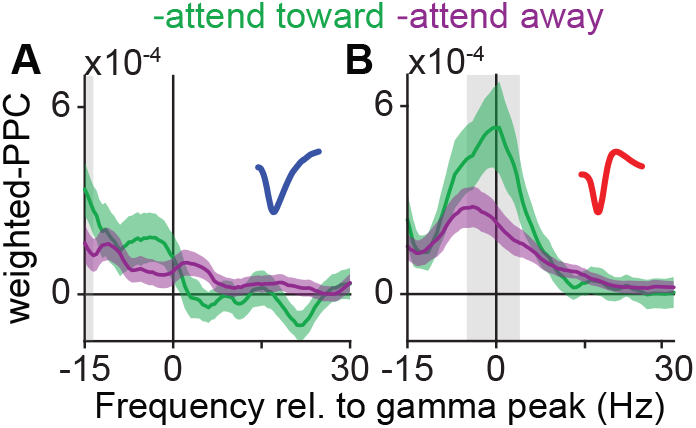
Cell-type specific effects on gamma-rhythmic synchronization persist after controlling for SNR. **(A,B)** Same as Fig. 2E, but after weighting the PPC spectrum corresponding to each cell by the cell’s number of spikes.

**Fig. S4:**
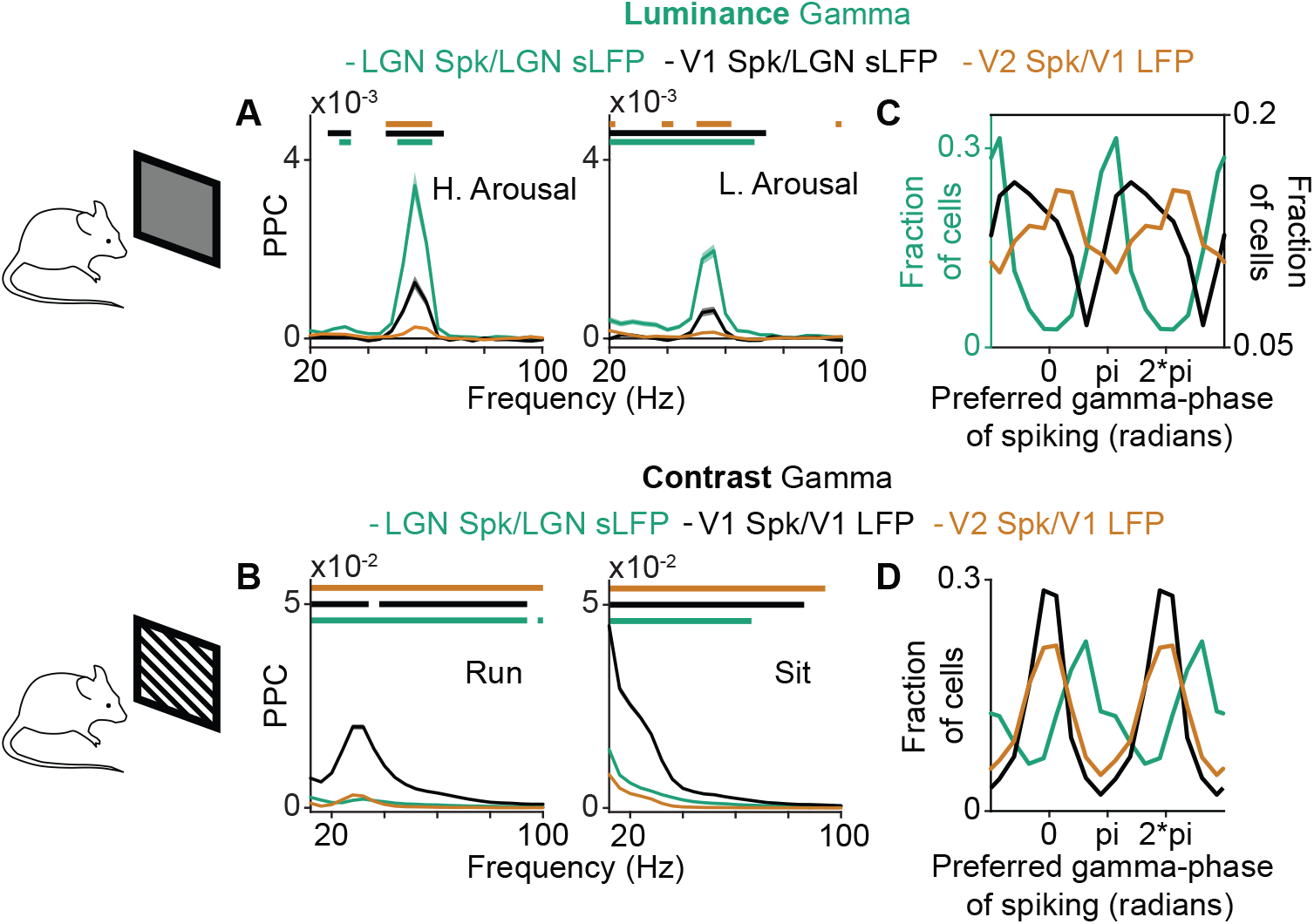
Feedforward gamma-band synchronization between LGN, V1, and V2 in the mouse, and the effect of behavioral state on its strength. **(A)** PPC between LGN sLFPs and LGN single unit (SU) spiking (*N* = 2215/2119), LGN sLFPs and V1 single unit (SU) spiking (*N* = 2700/2917), or V1 LFPs and V2 single unit (SU) spiking (*N* = 11962/13121), under the luminance gamma condition. Left: Analyses based on periods of high arousal. Right: Analyses based on periods of low arousal. Green, black, and orange horizontal bars in PPC spectra designate significantly different frequency bins between the spectra of LGN SUs and V1 SUs, between the spectra of LGN SUs and V2 SUs, and between the spectra of V1 SUs and V2 SUs, respectively. **(B)** Same as A, but between V1 LFPs and LGN single unit (SU) spiking (*N* = 2001/2109), V1 LFPs and V1 single unit (SU) spiking (*N* = 2169/2435), or V1 LFPs and V2 single unit (SU) spiking (*N* = 7463/8387), under the grating gamma condition. Left: Analyses based on periods when the animal ran. Right: Analyses based on periods when the animal was stationary. **(A,B)** Confidence intervals designate the SEM across cells (gray rectangles designate significantly different frequency bins between populations, randomization test across cells, FDR correction for multiple comparisons with a threshold of pĵ0.05). **(C)** Mean LGN-gamma phase of spiking for LGN (green, left y-axis) or V1 (black, right y-axis) SUs, and mean V1-gamma phase of spiking for V2 SUs (orange, right y-axis) (mean phase difference between populations = 1.51 radians, randomization test across relative phases, p = 0), under the condition of luminance-gamma. **(D)** Mean V1-gamma phase of spiking for LGN (green), V1 (black) or V2 SUs (orange) (mean phase difference between populations = 1.51 radians, randomization test across relative phases, p = 0), under the condition of grating-gamma.

**Fig. S5:**
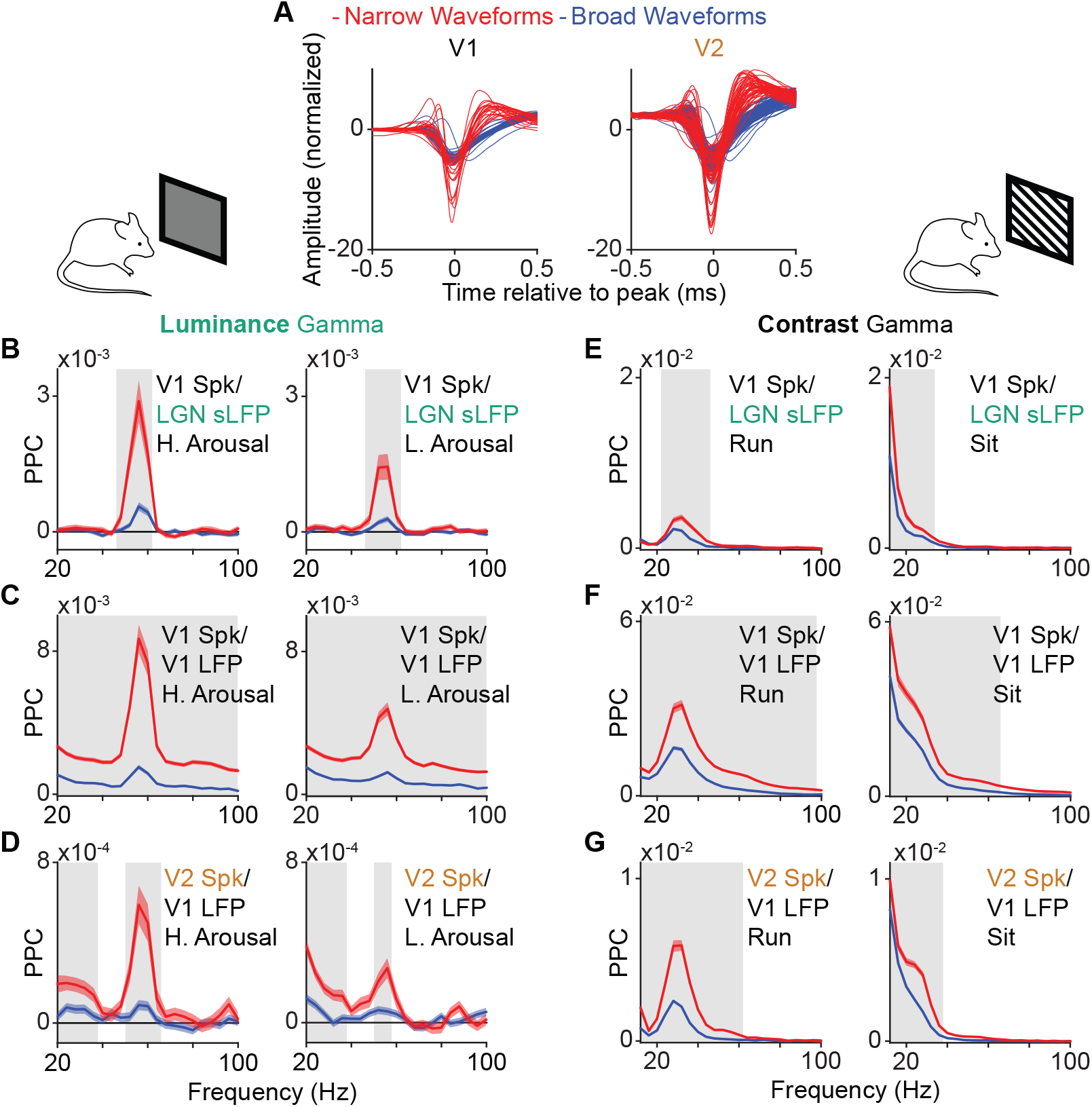
The strength of intra- and inter-areal gamma-band synchronization for different cell types depends on behavioral state in the mouse. **(A)** Same as Fig. 2A but for cells in mouse V1 (left) and V2 (right), for an example session. **(B)** Same as Fig. 2B but for LGN sLFPs and V1 SUs, under the condition of luminance gamma (BW: *N* = 1913/2113, NW: *N* = 698/718). **(C)** Same as Fig. 2B but for V1 LFPs and V1 SUs, under the condition of luminance gamma (BW: *N* = 2625/2950, NW: *N* = 928/962). **(D)** Same as Fig. 2B but for V1 LFPs and V2 SUs, under the condition of luminance gamma (BW: *N* = 8844*/*9931, NW: *N* = 2571*/*2595). **(B-D)** Left: Analyses based on periods of high arousal. Right: Analyses based on periods of low arousal. **(E)** Same as Fig. 2B but for LGN sLFPs and V1 SUs, under the condition of grating gamma (BW: *N* = 1186*/*1435, NW *N* = 476*/*500). **(F)** Same as Fig. 2B but for V1 LFPs and V1 SUs, under the condition of grating gamma (BW: *N* = 1519*/*1747, NW: *N* = 582*/*615). **(G)** Same as Fig. 2B but for V1 LFPs and V2 SUs, under the condition of grating gamma (BW: *N* = 5477*/*6091, NW *N* = 1585*/*1638). **(E-G)** Left: Analyses based on periods when the animal ran. Right: Analyses based on periods when the animal was stationary.

**Fig. S6:**
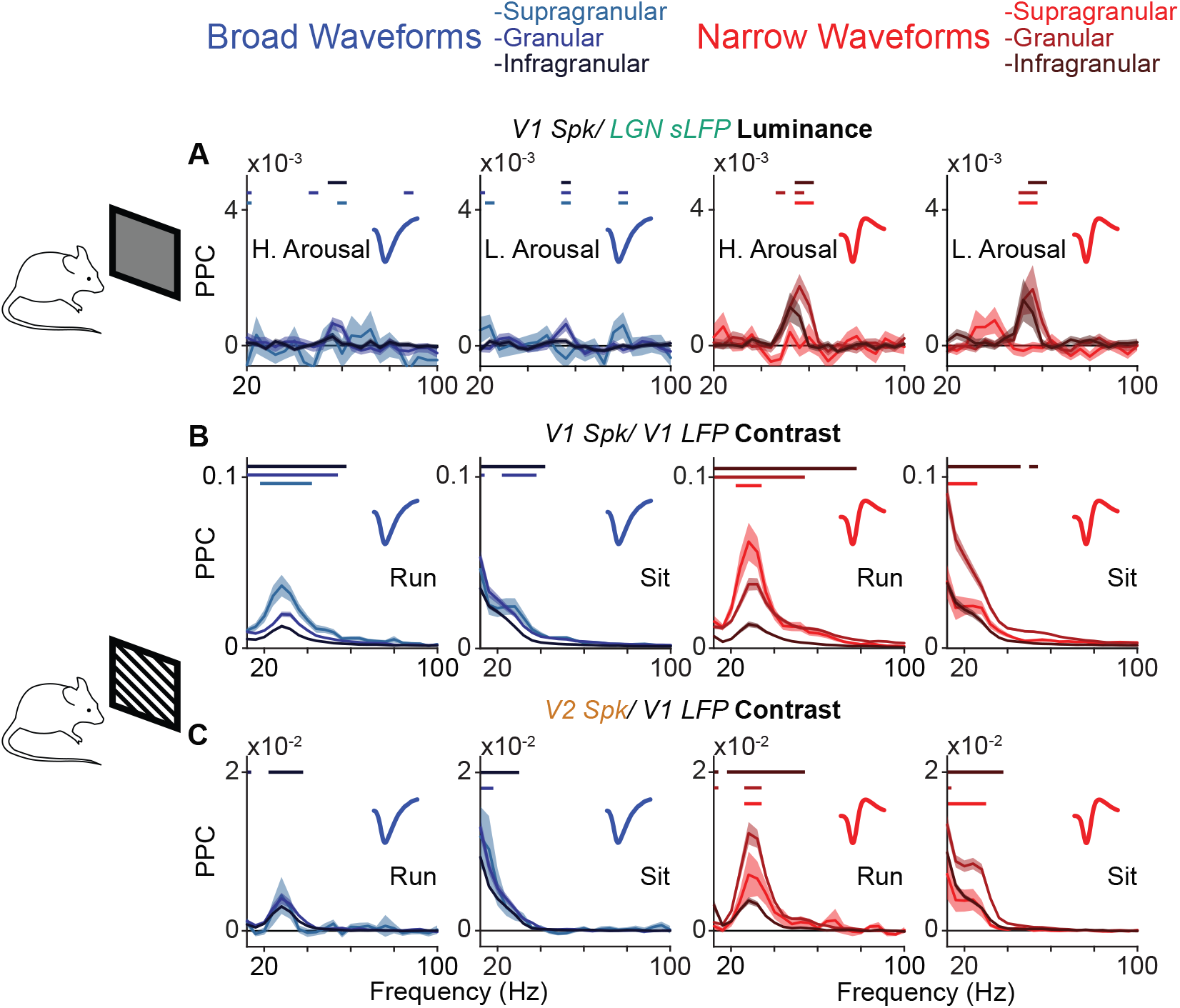
Laminar patterns in the effect of behavioral state on intra- and inter-areal gamma-band synchronization in the mouse. **(A)** Same as Fig. 3E but for LGN sLFPs and V1 SUs, under the condition of luminance gamma (BW: *Nsup*. = 50/110, *Ngra*. = 325/375, *Ninf*. = 769/774, NW: *Nsup*. = 41/44, *Ngra*. = 197/196, *Ninf*. = 265/259). **(B)** Same as Fig. 3E but for V1 LFPs and V1 SUs, under the condition of grating gamma (BW: *Nsup*. = 31/32, *Ngra*. = 154/156, *Ninf*. = 197/204, NW: *Nsup*. = 23/55, *Ngra*. = 232*/*282, *Ninf*. = 556/626). **(C)** Same as Fig. 3E but for V2 LFPs and V1 SUs, under the condition of grating gamma (BW: *Nsup*. = 23/41, *Ngra*. = 428/550, *Ninf*. = 2187/2449, NW: *Nsup*. = 11/15, *Ngra*. = 281/282, *Ninf*. = 544/571). **(A-C)** First and third panels from the left: Analyses based on periods when the animal ran. Second and fourth panels from the left: Analyses based on periods when the animal was stationary.

**Fig. S7:**
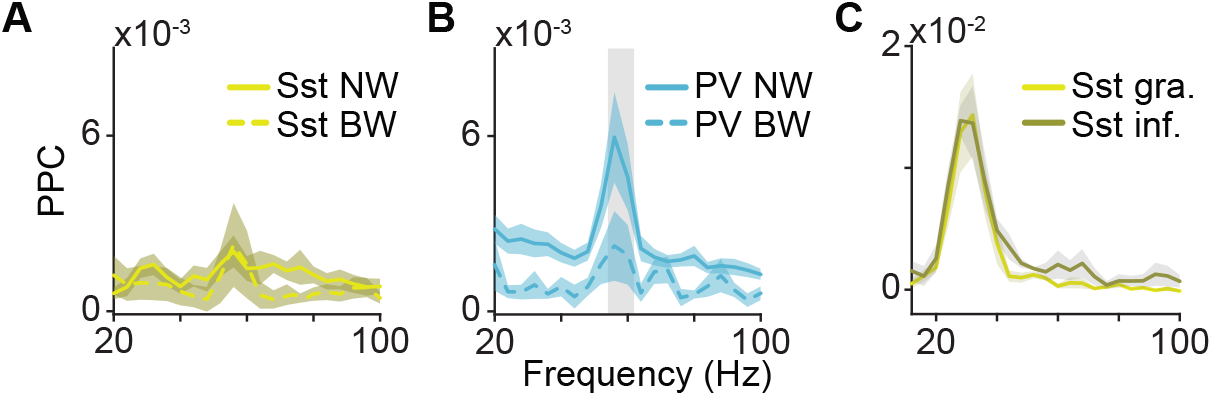
Inter-areal gamma band synchronization between V1 spiking and LGN sLFPs involve NW PV+ cells. **(A)** PPC between V1 LFPs and V1 PV+ cell spiking under the luminance-gamma condition, for BW (*N* = 19) or NW cells (*N* = 10). **(B)** Mean PPC spectrum between V1 LFPs and V1 PV+ cell spiking under the luminance-gamma condition, for BW (*N* = 9) or NW cells (*N* = 38). **(C)** Mean PPC spectrum between V1 LFPs and V2 Sst+ cell spiking under the luminance-gamma condition, located in the granular layer (*N* = 2) or the infragranular layers (*N* = 2). **(A-C)** Statistical comparisons are done in the same way as in Fig. 2B.

## References

Akam, T.E., Kullmann, D.M., 2012. Efficient “Communication through Coherence” requires oscillations structured to minimize interference between signals. PLOS Comp. Biol. 8, e1002760. doi:10.1371/journal.pcbi.1002760.

Atallah, B.V., Bruns, W., Carandini, M., Scanziani, M., 2012. Parvalbumin-expressing interneurons linearly transform cortical responses to visual stimuli. Neuron 73, 159–170.

Batista-Brito, R., Zagha, E., Ratliff, J.M., Vinck, M., 2018. Modulation of cortical circuits by top-down processing and arousal state in health and disease. Current opinion in neurobiology 52, 172–181.

Beaulieu-Laroche, L., Toloza, E.H., Van der Goes, M.S., Lafourcade, M., Barnagian, D., Williams, Z.M., Eskandar, E.N., Frosch, M.P., Cash, S.S., Harnett, M.T., 2018. Enhanced dendritic compartmentalization in human cortical neurons. Cell 175, 643–651.

Bernander, O., Douglas, R.J., Martin, K., Koch, C., 1991. Synaptic background activity influences spatiotemporal integration in single pyramidal cells. Proceedings of the National Academy of Sciences 88, 11569–11573.

Bernander, O., Koch, C., Usher, M., 1994. The effect of synchronized inputs at the single neuron level. Neural Computation 6, 622–641.

Bosman, C.A., Schoffelen, J.M., Brunet, N., Oostenveld, R., Bastos, A.M., Womelsdorf, T., Rubehn, B., Stieglitz, T., De Weerd, P., Fries, P., 2012. Attentional Stimulus Selection through Selective Synchronization between Monkey Visual Areas. Neuron 75, 875–888. doi:10.1016/j.neuron.2012.06.037.

Bruno, R.M., Sakmann, B., 2006. Cortex is driven by weak but synchronously active thalamocortical synapses. Science 312, 1622–1627.

Buffalo, E.A., Fries, P., Landman, R., Liang, H., Desimone, R., 2010. A backward progression of attentional effects in the ventral stream. Proc. Natl. Acad. Sci. U.S.A. 107, 361–365. doi:10.1073/pnas.0907658106.

Carandini, M., Heeger, D.J., 2012. Normalization as a canonical neural computation. Nature Reviews Neuroscience 13, 51–62.

Cardin, J.A., Carlén, M., Meletis, K., Knoblich, U., Zhang, F., Deisseroth, K., Tsai, L.H., Moore, C.I., 2009. Driving fast-spiking cells induces gamma rhythm and controls sensory responses. Nature 459, 663–667. doi:10.1038/nature08002.

Chaudhuri, R., Knoblauch, K., Gariel, M.A., Kennedy, H., Wang, X.J., 2015. A large-scale circuit mechanism for hierarchical dynamical processing in the primate cortex. Neuron 88, 419–431.

Dasilva, M., Brandt, C., Gotthardt, S., Gieselmann, M.A., Distler, C., Thiele, A., 2019. Cell class-specific modulation of attentional signals by acetylcholine in macaque frontal eye field. Proceedings of the National Academy of Sciences 116, 20180–20189.

Debes, S.R., Dragoi, V., 2023. Suppressing feedback signals to visual cortex abolishes attentional modulation. Science 379, 468–473.

Desimone, R., Duncan, J., et al., 1995. Neural mechanisms of selective visual attention. Annual review of neuroscience 18, 193–222.

Douglas, R.J., Martin, K.A., 2004. Neuronal circuits of the neocortex. Annu. Rev. Neurosci. 27, 419–451.

Felleman, D., C Van Essen, D., 1991. Felleman, D. J. & Van Essen, V. C. Distributed hierarchical processing in primate visual cortex. Cereb. Cortex 1, 1–47. volume 1. doi:10.1093/cercor/1.1.1.

Ferro, D., van Kempen, J., Boyd, M., Panzeri, S., Thiele, A., 2021a. Directed information exchange between cortical layers in macaque v1 and v4 and its modulation by selective attention. Proceedings of the National Academy of Sciences 118.

Ferro, D., van Kempen, J., Boyd, M., Panzeri, S., Thiele, A., 2021b. Directed information exchange between cortical layers in macaque v1 and v4 and its modulation by selective attention. Proceedings of the National Academy of Sciences 118, e2022097118.

Fries, P., 2005. A mechanism for cognitive dynamics: neuronal communication through neuronal coherence. Trends in cognitive sciences 9, 474–480.

Fries, P., 2015. Rhythm for Cognition: Communication Through Coherence. Neuron 88, 220–235. doi:10.1016/j.neuron.2015.09.034.Rhythms,arXiv:15334406.

Gentet, L.J., Kremer, Y., Taniguchi, H., Huang, Z.J., Staiger, J.F., Petersen, C.C., 2012. Unique functional properties of somatostatin-expressing gabaergic neurons in mouse barrel cortex. Nature neuroscience 15, 607–612.

Gieselmann, M., Thiele, A., 2008. Comparison of spatial integration and surround suppression characteristics in spiking activity and the local field potential in macaque v1. European Journal of Neuroscience 28, 447–459.

Gray, H., Bertrand, H., Mindus, C., Flecknell, P., Rowe, C., Thiele, A., 2016. Physiological, behavioral, and scientific impact of different fluid control protocols in the rhesus macaque (macaca mulatta). Eneuro 3.

Gregoriou, G.G., Rossi, A.F., Ungerleider, L.G., Desimone, R., 2014. Lesions of prefrontal cortex reduce attentional modulation of neuronal responses and synchrony in V4. Nat. Neurosci. 17, 1003–1011. doi:10.1038/nn.3742.

Grothe, I., Neitzel, S.D., Mandon, S., Kreiter, A.K., 2012. Switching Neuronal Inputs by Differential Modulations of Gamma-Band Phase-Coherence. J. Neurosci. 32, 16172–16180. doi:10.1523/JNEUROSCI.0890-12.2012.

Guzman, S.J., Schlögl, A., Espinoza, C., Zhang, X., Suter, B.A., Jonas, P., 2021. How connectivity rules and synaptic properties shape the efficacy of pattern separation in the entorhinal cortex– dentate gyrus–ca3 network. Nature Computational Science 1, 830–842.

Hamilton, L.S., Sohl-Dickstein, J., Huth, A.G., Carels, V.M., Deisseroth, K., Bao, S., 2013. Optogenetic activation of an inhibitory network enhances feedforward functional connectivity in auditory cortex. Neuron 80, 1066–1076.

Hermes, D., Petridou, N., Kay, K.N., Winawer, J., 2019. An image-computable model for the stimulus selectivity of gamma oscillations. Elife 8, e47035.

Izhikevich, E.M., Desai, N.S., Walcott, E.C., Hoppensteadt, F.C., 2003. Bursts as a unit of neural information: selective communication via resonance. Trends in neurosciences 26, 161–167.

van Kempen, J., Gieselmann, M.A., Boyd, M., Steinmetz, N.A., Moore, T., Engel, T.A., Thiele, A., 2021. Top-down coordination of local cortical state during selective attention. Neuron 109, 894–904.

Kim, H., Ahrlund-Richter, S., Wang, X., Deisseroth, K., Carlén, M., 2016. Prefrontal parvalbumin neurons in control of attention. Cell 164, 208–218.

Kreiter, A.K., 2006. How do we model attention-dependent signal routing? Neural networks 19, 1443–1444.

Markov, N.T., Vezoli, J., Chameau, P., Falchier, A., Quilodran, R., Huissoud, C., Lamy, C., Misery, P., Giroud, P., Ullman, S., Barone, P., Dehay, C., Knoblauch, K., Kennedy, H., 2014. Anatomy of hierarchy: Feedforward and feedback pathways in macaque visual cortex. Journal of Comparative Neurology 522, 225–259. doi:10.1002/cne.23458.

Maunsell, J.H., 2015. Neuronal mechanisms of visual attention. Annual review of vision science 1, 373.

McAdams, C.J., Maunsell, J.H., 1999. Effects of attention on orientation-tuning functions of single neurons in macaque cortical area v4. Journal of Neuroscience 19, 431–441.

McCormick, D.A., Connors, B.W., Lighthall, J.W., Prince, D.A., 1985. Comparative electrophysiology of pyramidal and sparsely spiny stellate neurons of the neocortex. Journal of neurophysiology 54, 782–806.

Mitchell, J.F., Sundberg, K.A., Reynolds, J.H., 2007. Differential Attention-Dependent Response Modulation across Cell Classes in Macaque Visual Area V4. Neuron 55, 131–141. doi:10.1016/j.neuron.2007.06.018.

Mitra, P.P., Pesaran, B., 1999. Analysis of dynamic brain imaging data. Biophysical journal 76, 691–708.

Mitzdorf, U., 1985. Current source-density method and application in cat cerebral cortex: investigation of evoked potentials and eeg phenomena. Physiological reviews 65, 37–100.

Montijn, J.S., Seignette, K., Howlett, M.H., Cazemier, J.L., Kamermans, M., Levelt, C.N., Heimel, J.A., 2021. A parameter-free statistical test for neuronal responsiveness. Elife 10, e71969.

Montijn, J.S., Vinck, M., Pennartz, C.M., 2014. Population coding in mouse visual cortex: response reliability and dissociability of stimulus tuning and noise correlation. Frontiers in computational neuroscience 8, 58.

Moradi Chameh, H., Rich, S., Wang, L., Chen, F.D., Zhang, L., Carlen, P.L., Tripathy, S.J., Valiante, T.A., 2021. Diversity amongst human cortical pyramidal neurons revealed via their sag currents and frequency preferences. Nature communications 12, 1–15.

Nichols, T.E., Holmes, A.P., 2002. Nonparametric permutation tests for functional neuroimaging: a primer with examples. Human brain mapping 15, 1–25.

Noudoost, B., Chang, M.H., Steinmetz, N.A., Moore, T., 2010. Topdown control of visual attention. Current Opinion in Neurobiology 20, 183–190. doi:10.1016/j.conb.2010.02.003.

Palmigiano, A., Geisel, T., Wolf, F., Battaglia, D., 2017. Flexible information routing by transient synchrony. Nature neuroscience 20, 1014.

van Pelt, S., Boomsma, D.I., Fries, P., 2012. Magnetoencephalography in twins reveals a strong genetic determination of the peak frequency of visually induced gamma-band synchronization. Journal of Neuroscience 32, 3388–3392.

Pesaran, B., Vinck, M., Einevoll, G., Sirota, A., Fries, P., Siegel, M., Truccolo, W., Schroeder, C., Srinivasan, R., 2018. Investigating large-scale brain dynamics using field potential recordings: analysis and interpretation. Nat. Neurosci..

Peter, A., Uran, C., Klon-Lipok, J., Roese, R., van Stijn, S., Barnes, W., Dowdall, J.R., Singer, W., Fries, P., Vinck, M., 2019a. Surface color and predictability determine contextual modulation of v1 firing and gamma oscillations. Elife 8, e42101.

Peter, A., Uran, C., Klon-Lipok, J., Roese, R., Van Stijn, S., Barnes, W., Dowdall, J.R., Singer, W., Fries, P., Vinck, M., 2019b. Surface color and predictability determine contextual modulation of V1 firing and gamma oscillations. eLife 8, e42101.

Pfeffer, C.K., Xue, M., He, M., Huang, Z.J., Scanziani, M., 2013. Inhibition of inhibition in visual cortex: the logic of connections between molecularly distinct interneurons. Nature neuroscience 16, 1068–1076.

Pike, F.G., Goddard, R.S., Suckling, J.M., Ganter, P., Kasthuri, N., Paulsen, O., 2000. Distinct frequency preferences of different types of rat hippocampal neurones in response to oscillatory input currents. The Journal of Physiology 529, 205.

Ray, S., Maunsell, J.H., 2010. Differences in Gamma Frequencies across Visual Cortex Restrict Their Possible Use in Computation. Neuron 67, 885–896. doi:10.1016/j.neuron.2010.08.004.

Roberts, M.J., Lowet, E., Brunet, N.M., Ter Wal, M., Tiesinga, P., Fries, P., De Weerd, P., 2013. Robust Gamma Coherence between Macaque V1 and V2 by Dynamic Frequency Matching. Neuron 78, 523–536. doi:10.1016/j.neuron.2013.03.003.

Saleem, A.B., Lien, A.D., Krumin, M., Haider, B., Rosón, M.R., Ayaz, A., Reinhold, K., Busse, L., Carandini, M., Harris, K.D., 2017. Subcortical Source and Modulation of the Narrowband Gamma Oscillation in Mouse Visual Cortex. Neuron 93, 315–322. doi:10.1016/j.neuron.2016.12.028.

Salinas, E., Sejnowski, T.J., 2001. Correlated neuronal activity and the flow of neural information. Nature reviews neuroscience 2, 539–550.

Scala, F., Kobak, D., Shen, S., Bernaerts, Y., Laturnus, S., Cadwell, C., Hartmanis, L., Froudarakis, E., Castro, J., Tan, Z., Papadopoulos, S., Patel, S., Sandberg, R., Berens, P., Jiang, X., Tolias, A., 2019. Layer 4 of mouse neocortex differs in cell types and circuit organization between sensory areas. Nature Communications 10, 1–12. doi:10.1038/s41467-019-12058-z.

Schmitzer-Torbert, N., Jackson, J., Henze, D., Harris, K., Redish, A., 2005. Quantitative measures of cluster quality for use in extracellular recordings. Neuroscience 131, 1–11. doi:10.1016/j.neuroscience.2004.09.066.

Schneider, M., Broggini, A.C., Dann, B., Tzanou, A., Uran, C., Sheshadri, S., Scherberger, H., Vinck, M., 2021. A mechanism for inter-areal coherence through communication based on connectivity and oscillatory power. Neuron 109, 4050–4067.

Schomburg, E.W., Fernández-Ruiz, A., Mizuseki, K., Berényi, A., Anastassiou, C.A., Koch, C., Buzsáki, G., 2014. Theta phase segregation of input-specific gamma patterns in entorhinal-hippocampal networks. Neuron 84, 470–485.

Senzai, Y., Fernandez-Ruiz, A., Buzsáki, G., 2019. Layer-specific physiological features and interlaminar interactions in the primary visual cortex of the mouse. Neuron 101, 500–513.

Shen, S., Jiang, X., Scala, F., Fu, J., Fahey, P., Kobak, D., Tan, Z., Zhou, N., Reimer, J., Sinz, F., et al., 2022. Distinct organization of two cortico-cortical feedback pathways. Nature Communications 13, 6389.

Siegle, J.H., Jia, X., Durand, S., Gale, S., Bennett, C., Graddis, N., Heller, G., Ramirez, T.K., Choi, H., Luviano, J.A., et al., 2021. Survey of spiking in the mouse visual system reveals functional hierarchy. Nature 592, 86–92.

Sohal, V.S., Zhang, F., Yizhar, O., Deisseroth, K., 2009. Parvalbumin neurons and gamma rhythms enhance cortical circuit performance. Nature 459, 698–702.

Thiele, A., Delicato, L.S., Roberts, M., Gieselmann, M.A., 2006. A novel electrode–pipette design for simultaneous recording of extracellular spikes and iontophoretic drug application in awake behaving monkeys. Journal of neuroscience methods 158, 207–211.

Uran, C., Peter, A., Lazar, A., Barnes, W., Klon-Lipok, J., Shapcott, K.A., Roese, R., Fries, P., Singer, W., Vinck, M., 2022. Predictive coding of natural images by v1 firing rates and rhythmic synchronization. Neuron 110, 1240–1257.

Vaidya, S.P., Johnston, D., 2013. Temporal synchrony and gamma-to-theta power conversion in the dendrites of ca1 pyramidal neurons. Nature neuroscience 16, 1812–1820.

Van Kerkoerle, T., Self, M.W., Roelfsema, P.R., 2017. Layer-specificity in the effects of attention and working memory on activity in primary visual cortex. Nature communications 8, 13804.

Veit, J., Hakim, R., Jadi, M.P., Sejnowski, T.J., Adesnik, H., 2017. Cortical gamma band synchronization through somatostatin interneurons. Nat. Neurosci 20, 951.

Vezoli, J., Magrou, L., Goebel, R., Wang, X.J., Knoblauch, K., Vinck, M., Kennedy, H., 2021a. Cortical hierarchy, dual counterstream architecture and the importance of top-down generative networks. Neuroimage 225, 117479.

Vezoli, J., Magrou, L., Goebel, R., Wang, X.J., Knoblauch, K., Vinck, M., Kennedy, H., 2021b. Cortical hierarchy, dual counterstream architecture and the importance of top-down generative networks. Neuroimage 225, 117479.

Vigneswaran, G., Kraskov, A., Lemon, R.N., 2011. Large identified pyramidal cells in macaque motor and premotor cortex exhibit “thin spikes”: implications for cell type classification. Journal of Neuroscience 31, 14235–14242.

Vinck, M., Battaglia, F.P., Womelsdorf, T., Pennartz, C., 2012. Improved measures of phase-coupling between spikes and the local field potential. Journal of computational neuroscience 33, 53–75.

Vinck, M., Womelsdorf, T., Buffalo, E.A., Desimone, R., Fries, P., 2013. Attentional Modulation of Cell-Class-Specific Gamma-Band Synchronization in Awake Monkey Area V4. Neuron 80, 1077–1089. doi:10.1016/j.neuron.2013.08.019.

Wilson, N.R., Runyan, C.A., Wang, F.L., Sur, M., 2012. Division and subtraction by distinct cortical inhibitory networks in vivo. Nature 488, 343–348.

Xu, H., Jeong, H.Y., Tremblay, R., Rudy, B., 2013. Neocortical somatostatin-expressing gabaergic interneurons disinhibit the thalamorecipient layer 4. Neuron 77, 155–167.

Zhang, S., Xu, M., Kamigaki, T., Do, J.P.H., Chang, W.C., Jenvay, S., Miyamichi, K., Luo, L., Dan, Y., 2014. Long-range and local circuits for top-down modulation of visual cortex processing. Science 345, 660–665. doi:10.1126/science.1254126.

